# A systematic review of laboratory investigations into the pathogenesis of avian influenza viruses in wild avifauna of North America

**DOI:** 10.1101/2024.05.06.592734

**Authors:** Matthew Gonnerman, Christina Leyson, Jeffery D. Sullivan, Mary J. Pantin-Jackwood, Erica Spackman, Jennifer M. Mullinax, Diann J. Prosser

## Abstract

The lack of consolidated information regarding wild bird species’ response to avian influenza virus (AIV) infection is a challenge for conservation managers, researchers, and related sectors such as public health and commercial poultry. Such information could be used to model complex disease dynamics within communities, prioritize species for surveillance and conservation efforts, or identify species more likely to facilitate spillover into domestic animals or humans. Using two independent searches, we reviewed published literature for studies describing wild bird species experimentally infected with avian influenza to assess host species’ relative susceptibility to AIVs. Additionally, we summarized broad-scale parameters for elements such as shedding duration and minimum infectious dose that can be used in transmission modeling efforts. Our synthesis documented that waterfowl (i.e., Anatidae) comprise the vast majority of published AIV pathobiology studies, whereas gulls and passerines were less represented in research despite evidence that they also are susceptible and contribute to high pathogenicity avian influenza disease dynamics. This study represents the first comprehensive effort to compile available literature regarding the pathobiology of AIV’s in all wild birds in over a decade. This database provides an opportunity to critically examine and assess what is known and identify where further insight is needed.

## INTRODUCTION

While low pathogenic avian influenza (LPAI) virus circulates naturally with minimal clinical signs in waterbirds in the orders Anseriformes and Charadriiformes [1,2], high pathogenicity avian influenza (HPAI) virus historically emerges following mutation or reassortments gained by LPAI viruses circulating in gallinaceous poultry [3], often originally introduced by wild waterfowl (typically Anatidae). The recently circulating and expanding HPAI H5 viruses of the Goose/Guangdong/1/1996 (Gs/GD) lineage are unusual because they have adapted to be carried by certain wild waterfowl species, thereby expanding exposure to poultry, wild birds, and more recently some mammals [4,5]. The original H5 Gs/GD virus was isolated in 1996 in southern China [6] and by 2013 had evolved into clade 2.3.4.4 with a notably improved ability to infect wild waterfowl including species that are generally not carriers of LPAI viruses [7,8]. In October 2020, the global reach of H5 Gs/GD viruses was again highlighted when clade 2.3.4.4b viruses emerged, before spreading to over 80 countries in the following years [5]. This current outbreak has been especially notable for reaching new hemispheres as it moved into North America in 2021 [9], South America in late 2022 [10] and the Antarctic region in 2023 [11], causing extreme impact on numerous populations of wild birds such as the Peruvian Pelican (*Pelecanus thagus*), Sandwich Tern (*Thalasseus sandvicensis*), and Common Crane (*Grus grus*) [4].

During this recent panzootic of HPAI, the conservation community was challenged to anticipate how diverse species might respond when exposed to novel viral strains. For instance, there has been great concern about potential impacts on penguin colonies and condor populations should AIV be introduced [12] and recent outbreaks in Antarctica and South America indicate the vulnerability of these species [11,12]. However, an extensive body of work has been produced by researchers focused on challenge studies with wild bird species in a laboratory setting with both LPAI and HPAI to determine factors including infectiousness, severity of clinical signs, and recovery potential [13,14]. However, given the limited resources available, such work has not encompassed the broad spectrum of potential species and viral strains [15]. This has left a substantial knowledge gap as AIVs continue to evolve and present risks for an ever-growing list of species [16]. Furthermore, the available literature is diffuse with no comprehensive review for wild species collated in nearly a decade [14,17].

The lack of consolidated information on the response of wild bird species to infection with AIV is an identified challenge to both conservation managers and researchers alike [16], with additional impacts on related sectors such as public health and commercial poultry [18,19]. For wildlife managers, it is important to understand the possible implications of an HPAI outbreak, especially for sensitive populations [4]. For instance, knowing if a population is at-risk to exposure and resilient to infection may play an important role in how that species is prioritized in allocation of resources for habitat protection, captive breeding programs, and other programs. Similarly, the diffuse knowledge base makes identifying data-informed parameters for transmission risk models more difficult [16,18]. While we can infer some information from wild surveillance and mortality data, such as which species have tested positive and which are most likely associated with mortality events, we cannot determine important metrics like length of infection or infectivity from these databases. The goal of this study is to collate all previous pathobiology studies on wild avifauna in North America, providing insight into current gaps in the available literature that may be filled by future targeted research. Additionally, we aim to develop broad scale parameters for elements such as shedding duration and minimum infectious dose that can be used in transmission modeling efforts until species and strain specific results become available.

## METHODS

### Literature Search

Using two independent searches, we reviewed the published literature for studies in which wild bird species were experimentally infected with avian influenza to assess the host species’ relative susceptibility to AIVs and ability to shed viruses. To provide reasonable bounds on which species we included in our search, we created a list of species from those reported to the U.S. Department of Agriculture Animal and Plant Health Inspection Service (USDA-APHIS) database [20] and the U.S. National Institute of Allergy and Infectious Diseases (NIAID)-sponsored Bioinformatics Resource Center Influenza Research Database (IRD) [21,22] as of Oct 16, 2023. We then supplemented this initial list with species that were both a high-priority conservation concern and at high risk of exposure to HPAI due to either shared habitat with or a close genetic relationship to known susceptible species. We conducted an initial search using Google Scholar during October and November of 2023 using the search criteria ‘“avian influenza” AND “[species name]” AND “challenge study” OR “experimental infection”‘. The search was conducted twice for each species of interest, where [species name] was replaced with either the common or scientific name. This initial search was intended to produce a representative, but not exhaustive, sample of research available for use in modeling avian influenza disease dynamics. As some species produced a disproportionately large number of papers (+1000) compared to the average (0–50), we limited our search to the first 5 pages of Google Scholar results to balance search time and capturing relevant information for all species.

While limiting search efforts likely contributed to missing papers, there are also concerns that Google Scholar is not an appropriate method for reproducible reviews of scientific literature [23,24]. So, after completion of this initial search, we identified several relevant studies providing information on unrepresented HPAI lineages not identified by Google Scholar by conducting a second systematic review using Web of Science. We expanded our search criteria as using the same search terms as the original review proved too restrictive when applied to Web of Science. We chose to simplify phrasing, remove quotations, and include additional language previously used in reference to challenge studies, resulting in a search criteria of ‘(TS=(“avian influenza” AND “[species name]”) AND TS=(challenge OR experiment OR characterization OR pathogenesis))’. We conducted the search as initially described for our first review, repeating the search for each common and scientific name for birds on our search list.

### Identifying Relevant Studies

Articles reviewed herein were considered for further review if the title, abstract, or methods discussed AIV challenge studies for wild bird species that range within North America. We excluded information for non-avian influenza viruses (i.e., mammal-derived strains of AIV) as well as information related to domestic and hybrid host bird species. There were multiple instances when a study challenged an individual with multiple viral strains in succession. In such cases, we recorded the viral subtype of the previous infection but otherwise, recorded this information as usual, as many wild animals are likely to have previous AIV infections. However, several studies examined the impact of vaccines or viral inhibitors on host birds or local environments (i.e., water treatments), in which case we recorded these studies in the database but did not include them in any summaries.

For studies that met our above criteria, we recorded a unique entry for each combination of host bird species, virus subtype, and challenge method (e.g., direct inoculation, contact, meat-fed) used within a study. For each entry, we recorded information regarding the host animal species, family, and age at inoculation, as well as the viral subtype, origin, whether the virus was a HPAI or LPAI pathotype, and HA genetic lineage. AIV lineages for subtypes other than goose/Guangdong/1996 H5 were described based on the HA gene of the most closely related grouping classified by naming conventions utilized in previous reports or general geographic and species origin from which similar isolates have been collected. Subtype H5 isolates were classified by clade if from the goose/Guangdong/1996 lineage [25].

To describe the experimental design of each study, we recorded inoculation method, inoculation dose, and swabbing schedule, with any further details that differentiated experimental groups within a study provided in a notes column. For each entry, we recorded the number of individuals in the study, the number of individuals identified as infected with AIV (i.e., virus identified in swabs, blood, or organs), and the number of mortalities. We separately counted direct mortalities from illness versus euthanasia due to symptoms deemed indicative of imminent mortality. In some studies, individuals were euthanized (hereafter “culled”) at predetermined time points throughout the study to assess viral titers in organs and tissue. We recorded these individuals separately from the prior two mortality categories to adjust estimates of mortality rate due to AIV infection. Information on mortality and culling did not include individuals culled at the end of the study (i.e., survivors). If provided, or if it could be calculated from the provided mortality data, we recorded the mean death time in days.

### Summarizing Viral Response

Across studies, viral response is measured and reported in a variety of ways which limited comparison among studies. Identification of viruses in swab samples can be performed using virus isolation or real-time RT-PCR methods[26], which are reported as at least one of multiple metrics that are not all comparable. Virus titer levels in isolates are commonly quantified and reported as the log_10_ 50% egg infective dose per milliliter (EID_50_/mL); however, alternative units can be used that produce estimates that are not comparable between methods [27]. Real-time RT-PCR results are readily reported as a cycle threshold (Ct) value, but additional steps need to be conducted to produce a comparable equivalent to the EID_50_/mL obtained from viral isolation [28]. As such, we only used information measured as EID_50_/mL from viral isolation or, if not available, EID_50_/mL equivalents from RT-PCR in our summaries. For simplicity, we hereafter use “shedding” to refer to the greatest daily-average viral titers identified in swabs. Information on shedding could be collected via swabbing the oropharyngeal (OP) and/or the cloacal (CL) routes, which are important prerequisites of animal-to-animal and environmental HPAI transmission respectively [13]. As shedding patterns between these pathways may have important but different implications for virus transmission, we recorded this information separately. We recorded information on viral shedding by the host animal, however this information is assessed and summarized in multiple formats across studies that are not directly comparable. For example, viral shedding quantities and durations were reported using a broad range of summary statistics, including mean, median, and maximum values for a given experimental group as well as by individual, sometimes for multiple time points throughout an infection. Thus, to describe the average capacity a host animal has for spreading the virus, we report the greatest mean daily shedding quantities for a given experimental group over the course of a study (Figure S1).

For all entries, we recorded the mean daily shedding quantities for a given experimental group and mean bird infectious dose (units in log_10_ 50% bird infectious doses per milliliter; BID_50_) if estimated. We also recorded shedding duration (days that an animal shed the virus), either as a single duration estimate where provided, or as a series of time points describing when the virus was first detected (D_1_; Figure S1), which day the greatest mean shedding rate occurred (D_Max_), the last day that the virus was detected in swabs (D_L_), and the next day animals were swabbed and virus was not detected following completion of shedding (D_Neg_). From this information and information on when swabbing occurred, we can calculate a shedding duration as the (D_L_ + D_Neg_)/2 – D_1_. In multiple instances, we encountered studies where shedding information was presented only in plot form. To interpret information within these papers, we used a web-based figure reader (PlotDigitizer; plotdigitizer.com) which is a free option for such data mining from scatter and line plots [29] and recorded its use.

Tables including entries for all inoculation doses, inoculation methods, and prior infection statuses were generated. To produce a final set of directly comparable metrics describing a species’ viral response, we subset our dataset to entries where animals were directly inoculated with a Gs/GD lineage HPAI of dose greater than 4 log_10_ EID_50_, which is a cutoff that consistently resulted in infection in most studies that estimated infectious dose.

Following the course of a successful infection of challenged individuals, we can further describe immune response for both animals that recovered and those that succumb to the disease. From surviving animals or animals euthanized during the course of recovering from infection, antibody production can be assessed via serological assays such as bELISA [30] to determine the strength and length of antibody response. In our review, we recorded whether a study collected serological information to describe antibody production following infection, as well as the maximum number of individuals recorded as positive for antibodies at a given timepoint. We can further study the severity and location of infection through necropsies of infected individuals, either to identify antigens or histological lesions in tissue samples [31,32]. If only a local infection has occurred, antigens or lesions will be isolated to the respiratory or digestive tract of the animal, whereas systematic infections of HPAI viruses can be identified throughout the body, indicating a more severe infection. Unfortunately, which organs were sampled varied amongst studies and studies were inconsistent in how they reported data (i.e., ranging from individual-level data to study-level summaries). Thus, we only present whether histopathology information was collected, and whether a group of animals had any individuals with local (i.e., lungs, trachea, digestive tract only) or systematic (i.e., any other organs) infections, identified by either lesions or antigens in a tissue sample. We note that immune response will be temporally variable relative to exposure as when any given individual was sampled will dramatically impact results.

### Comparison to Incidence of Avian Influenza in Wild Birds

To identify potential knowledge gaps in AIV research, we compared our review of the published research to the observed incidence of avian influenza in wild birds. Information on wild bird infections was obtained via the USDA-APHIS database [20] and accessed on Nov 16, 2023. The database provided information on encounter location, encounter date, viral subtype, host species, and how the animal was reported, and was summarized to the number of positives per species.

## RESULTS

### Summary of Relevant Studies

From our review database [52], 132 unique challenge studies for AIV in wild bird species were identified, and of these, 104 had at least one focal species from our list (Supplemental 1, Figure S2). Of these 104 studies, Anatidae was the most well-represented family identified (78 studies), followed by Laridae (11), Passeridae (8), Columbidae (6), and Scolopacidae (4; Figure 1, Table S1). Within Anatidae (Table S2), challenge studies were distributed across 18 species including mallard (*Anas platyrhynchos*; 54 studies), Muscovy duck (*Cairina moschata*; 11), Canada goose (*Branta candadensis*; 5), northern pintail (*Anas acuta*; 4), and wood duck (*Aix sponsa*; 4). Outside of Anatidae, house sparrow (*Passer domesticus*; 8), laughing gull (*Leucophaeus atricilla*; 7), and rock pigeon (*Columba livia*; 6) were the most studied. Of note, 6 of the top 10 families with the most reported AIV infections had no studies identified within our review (Figure 2), including Accipitridae (727 reported infections), Cathartidae (691), Strigidae (248), Pelecanidae (119), Corvidae (118), and Phalacrocoracidae (39). Forty-three studies challenged animals with LPAI subtypes and 74 with HPAI subtypes, of which 66 included viruses of the Gs/GD HPAI lineage (Figure 1). Only 5 of the 1282 animals challenged with LPAI in the identified studies perished over the course of the infection. Animals challenged with Gs/GD perished with greater frequency (67%, N = 1755) compared to those challenged with other lineages of HPAI (6%, N = 300). Information on viral susceptibility and shedding was only available from one species for 6 of the 10 families identified in the review, and for 3 of these families only 1 study was found (Table S3-S5). One of these, Ardeidae, had challenge data for only 3 individuals, of which 1 was infected and died.

**Figure 1.**
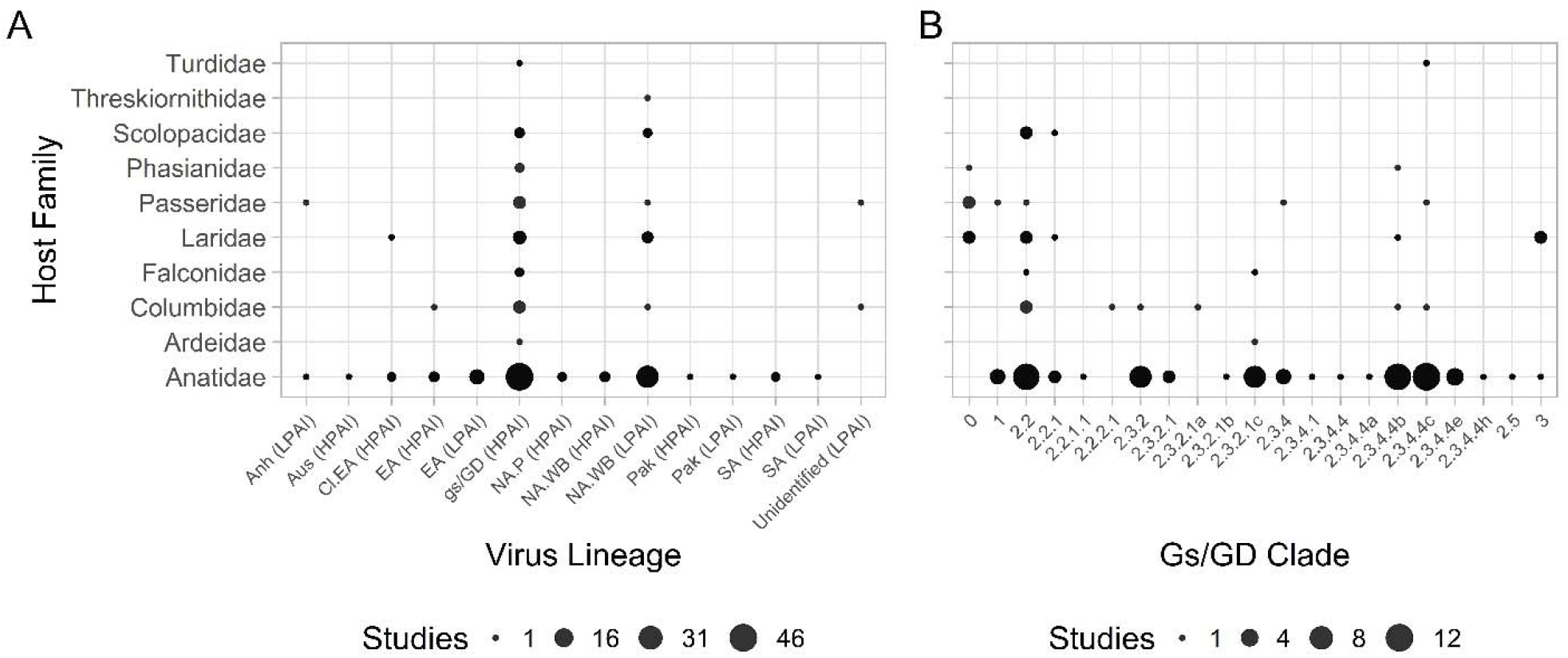
Bubble plot depicting the number of avian influenza virus challenge studies by viral lineage. We present information A) for all HPAI and LPAI lineages. Data is grouped by host species family and viral lineage and B) by clade of HPAI H5 viruses of the Goose/Guangdong/1/1996 (Gs/GD) lineage. Viral lineages identified include Anhui (Anh), Australian (Aus), Classical Eurasian (Cl.EA), Eurasian (EA), Gs/GD, North American poultry (NA.P), North American wild birds (NA.WB), Pakistan H7 (Pak), and South American (SA).

**Figure 2.**
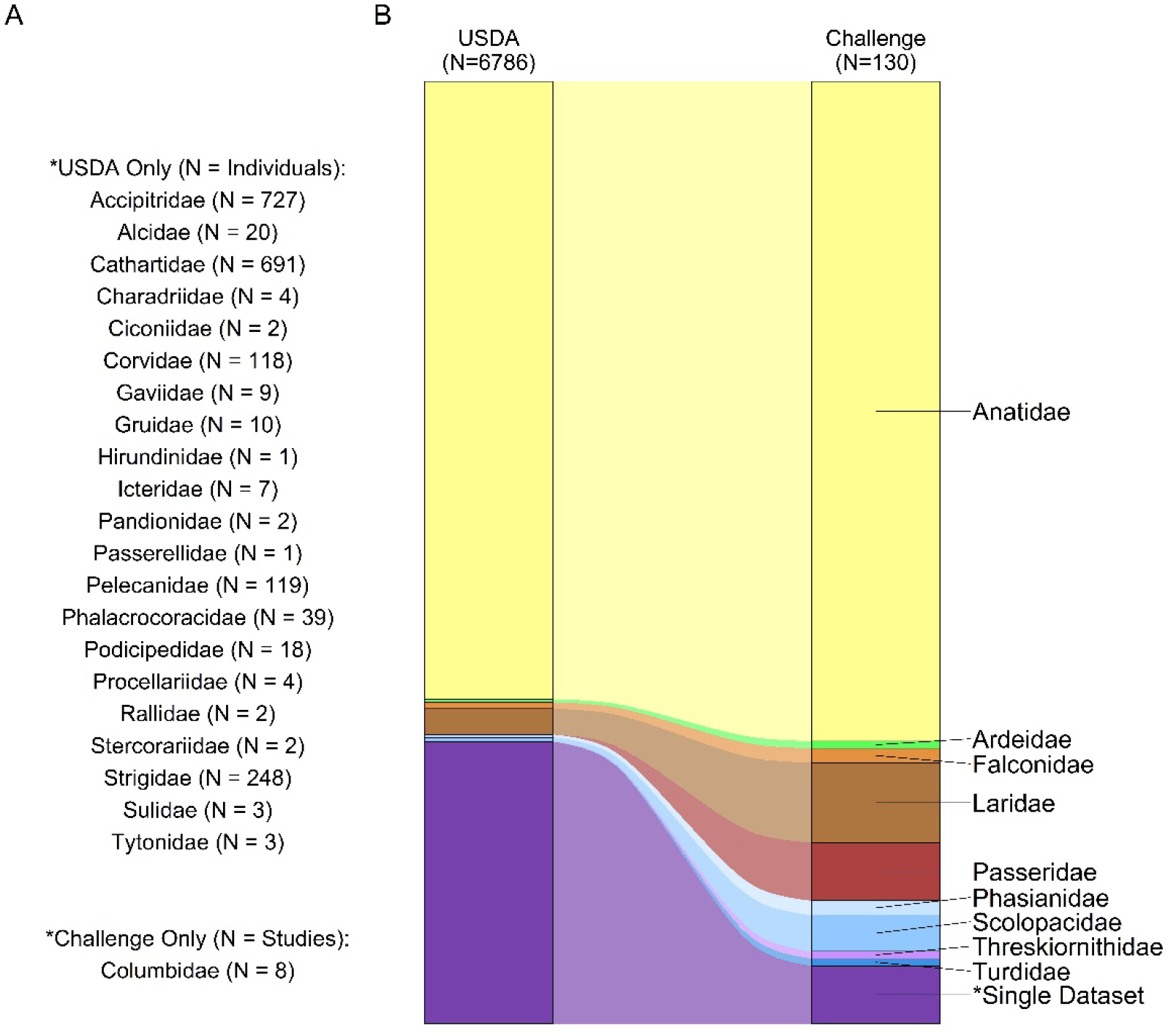
Comparison of the proportional composition of families identified in HPAI infections reported to the USDA-APHIS database [20] (accessed Nov 16, 2023) and HPAI challenge studies identified in our review. Families represented in only one dataset are listed with their respective sample size. We present A) a list of families which were only identified in one of the two datasets (i.e., “Single Dataset” in B) and B) a Sankey plot comparing proportional composition of families identified in each dataset.

### Summarizing Viral Response

Articles were subsetted according to challenge type (i.e., directly inoculated seronegative animals with >4 log_10_ EID_50_/mL of a Gs/GD HPAI viruses) then summarized (Figure 3, Table S3,S4). The resulting dataset contained information on only 24 species, of which 13 are from the Anatidae family. Within Anatidae, American black duck (*Anas rubripes*), blue-winged teal (*Spatula discors*), lesser scaup (*Aythya affinis*), Northern pintail, and redhead (*Aythya americana*) all exhibited 0% mortality when challenged with Gs/GD whereas Canada goose, mute swan (*Cygnus olor*), and trumpeter swan (*Cygnus buccinator*) all experienced 100% mortality. For waterfowl, oral shedding (measured by RNA presence from OP swabs) rates averaged 4.54 log_10_ EID_50_ and ranged from 1.1 log_10_ EID_50_ to 7.7 EID_50_ while cloacal shedding (measured by RNA presence from cloacal swabs (CL)) rates averaged 3.28 log_10_ EID_50_ and ranged from 0.9 log_10_ EID_50_ to 6.8 log_10_ EID_50_. Shedding durations averaged 5.72 and 5.21 days for oral and cloacal routes respectively. Trumpeter and mute swans exhibited the highest average oral (6.14 log_10_ EID_50_) and cloacal (4.46 log_10_ EID_50_) shedding rates respectively. The top eight oral shedders and seven of the top eight cloacal shedders were all from the Anatidae family. A number of these species, such as mallards and surf scoter (*Melanitta perspicillata*), also exhibited relatively low mortality rates (31% and 22 %) when challenged with an HPAI virus. Mallards exhibited higher than average shedding rates (4.78 log_10_ EID_50_ via OP, 3.52 log_10_ EID_50_ via CL) and durations (6.42 days via OP, 5.83 days via CL) across studies.

**Figure 3.**
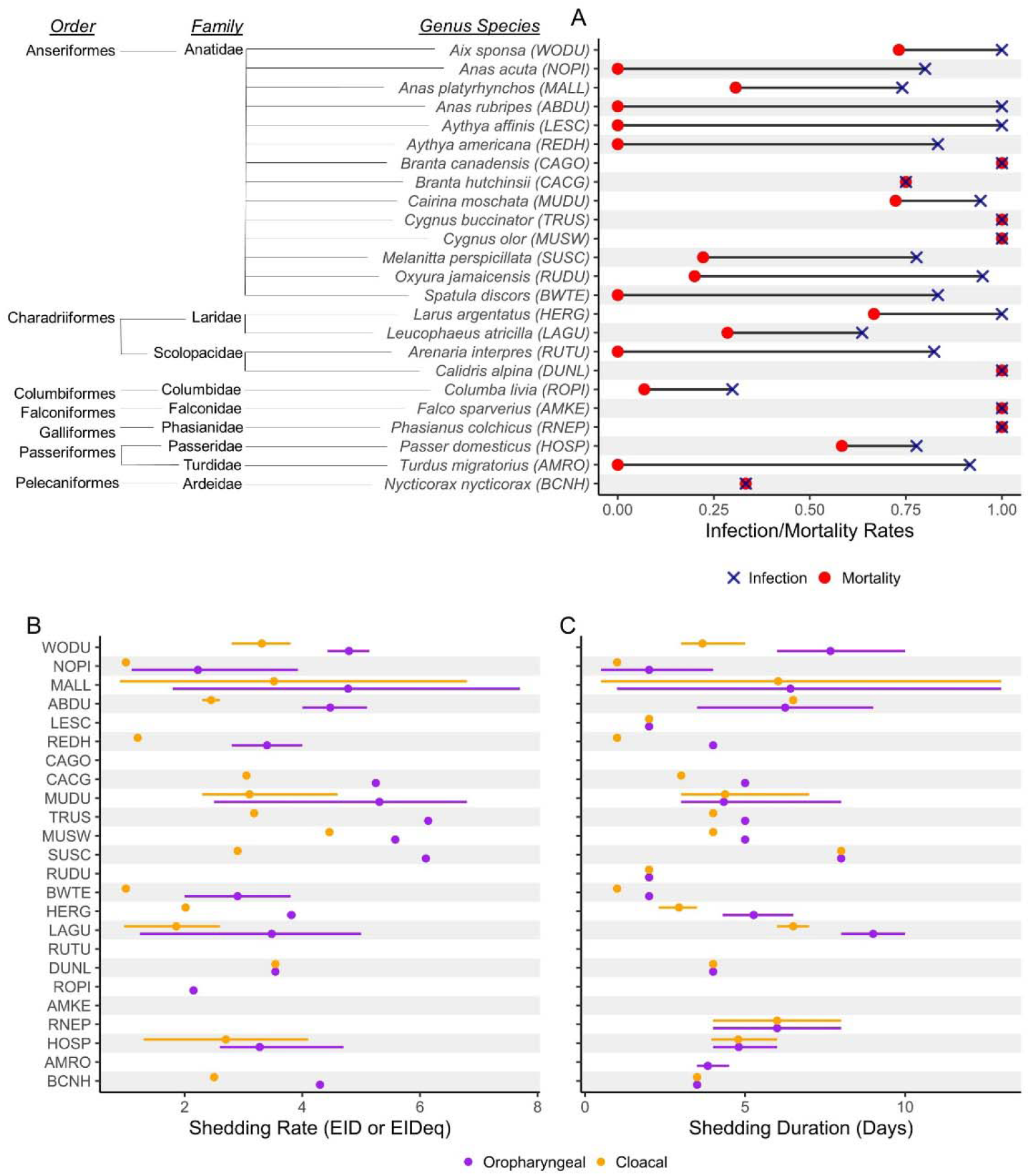
Disease response of seronegative host animals inoculated with >4 log_10_ EID_50_/mL of a Gs/GD HPAI. Across studies, we summarize and present A) infection (blue) and mortality (red) rates, B) shedding rate in log_10_ EID_50_ or EIDeq, and C) duration of shedding in days for oropharyngeal (purple) and cloacal (orange) routes. Abbreviations by common name, repeated in order across panels: Wood Duck (WODU), Northern Pintail (NOPI), Mallard (MALL), American Black Duck (ABDU), Lesser Scaup (LESC), Redhead (REDH), Canada Goose (CAGO), Cackling Goose (CACG), Muscovy Duck (MUDU), Trumpeter Swan (TRUS), Mute Swan (MUSW), Surf Scoter (SUSC), Ruddy Duck (RUDU), Blue-winged Teal (BWTE), Herring Gull (HERG), Laughing Gull (LAGU), Ruddy Turnstone (RUTU), Dunlin (DUNL), Rock Pigeon (ROPI), American Kestrel (AMKE), Ringed-neck pheasant (RNEP), House Sparrow (HOSP), American Robin (AMRO), Black-crowned Night Heron (BCNH)

We identified only a limited number of comparable studies for non-waterfowl species. For example, within Laridae, we identified herring gull (*Larus argentatus*) and laughing gull as the only species with data available. Herring gull exhibited high infection rates (100%) and moderate mortality rates (67%) when challenged and shed more via the OP route (3.82 log_10_ EID_50_ for 5.27 days) than the cloaca (2.01 log_10_ EID_50_ for 2.93 days). Comparatively, laughing gull had lower infection (64%) and mortality (29%) rates, longer shedding durations (9 days and 6.5 days for OP and CL), but similar shedding rates (3.48 log_10_ EID_50_ and 1.89 log_10_ EID_50_). Considering passerines, house sparrows had relatively high infection (78%) and mortality (58%) rates and shed moderately via both the OP and CL routes (3.28 log_10_ EID_50_ via OP, 2.7 log_10_ EID_50_ via CL) for approximately 4.8 days. Limited information was also available for American robins (*Turdus migratorius*), which were readily infected (92%) but showed no indication of negative impacts to survival (0% mortality). Rock pigeons, a similarly broadly abundant bird species, were much less readily infected (30%) and had similarly low mortality rates (7%). Within Falconidae, two studies challenged a total of 22 American kestrels (*Falco sparverius*), of which all perished.

When the dataset was expanded to include any animal challenged with any dose of Gs/GD HPAI (Table S5), we found Canada goose exhibited the greatest average viral titers in cloacal swabs (5.75 log_10_ EID_50_/mL, range 3.8-7.3) and trumpeter swan in OP swabs (6.14 log_10_ EID_50_/mL).

Laughing gulls had the longest observed shedding duration via the OP route (9 days post challenge, range 8-10), whereas surf scoters shed the longest via the CL route (8 days post challenge).

## DISCUSSION

This study represents the first comprehensive effort to compile available literature regarding the pathobiology of AIVs in all wild birds of North America in over a decade [13]. The database produced through this effort can serve as a tool for all researchers, providing generalized estimates of pathobiology parameters such as mortality and viral shedding for a variety of wild avian families. This database also provides the opportunity to critically examine previous research to assess what has been learned from these efforts as well as to identify where additional targeted efforts can provide further insight. However, we acknowledge the limitations of this resource, as the variability in methods used to assess viral response amongst studies limits our ability to compare among them. It should also be noted that infections in wild versus laboratory settings can produce divergent results [33], which should motivate further research when encountered. Thus we urge users to be thoughtful in their application of this database, and for future research to be exhaustive in their reporting and consistent with current common practices.

Our synthesis shows that waterfowl (i.e., Anatidae) comprise the vast majority of published AIV pathobiology studies. Such an emphasis is warranted given the widely recognized role of these species in the spread of AIVs [18,34]. Indeed, previous efforts have demonstrated that *Anatidae* are competent vectors for avian influenza viruses due to their high shedding rates [35,36], long duration of infection [37,38], and comparatively high prevalence in the wild [39,40]. However, while previous studies have provided us with valuable information on the general response to infection for Anseriformes, the importance of these species to the transmission of AIVs both across space [34,41] and across the wild waterfowl – domestic poultry interface [18,42] suggests that future work with these species could aim to develop better estimates of parameters needed for transmission risk modeling [43] such as minimum infectious dose which is currently only rarely included [44,45]. Similarly, the continued high representation of Anatidae in mortality events and surveillance sampling [20] emphasizes the importance of studies focused on including emerging viral strains to inform possible outcomes prior to outbreak events and to maintain current data.

There was a dramatic decrease in available research between Anseriformes and Laridae, the second most studied family group with 10 studies across 4 species. Laridae as a group had a high capacity for infection and shedding, consistent with the recognized importance of Laridae in the inter-continental spread and persistence of AIV [46,47]. Given the apparent susceptibility of these species in the current HPAI outbreak, their large geographic distribution, long-distance migrations, as well as frequent overlap with agriculture, Laridae seem to be prime candidates for continued pathobiology research [15]. For similar reasons, other sea birds, such as skuas [48], have been implicated as possible long-distance carriers of the virus, but a lack of available research limits our ability to infer potential dynamics.

While identified as having a role in the spatial movement of AIVs [49], predatory and scavenging species appear especially susceptible to infection and mortality based on data from the current outbreak [20], as well as limited pathobiology studies with species outside North America [50]. In fact, Accipitridae, Cathartidae, and Strigidae were three of the top four families with the most reported infections to USDA yet had no studies were identified by our review. However, Falconidae had among the highest infectious rates and mortality rates (100% for each), though this finding was based on a single species across two studies (the American Kestrel). Given the dramatic impacts of HPAI on these species there is a clear need to improve the understanding of all aspects of pathobiology of AIV in raptors and other predatory avian species. Additionally, efforts to model broader ecological transmission and persistence could benefit from studies that explore transmission through meat/carcass consumption instead of traditional inoculation [51].

Of course, many other species outside of those specifically mentioned above have been impacted by the ongoing outbreak of HPAI. Our summary demonstrates that seven families had over 100 reported HPAI infections to USDA [20], but only two of these families have been studied (Anatidae and Laridae). Of the 10 families identified in our review, only three reported challenge data from more than 1 species, and 1 of these only included three individuals in total. It should also be noted that these metrics rely on data from infections reported to USDA [20], so species that are under-sampled due to low detectability of sick or dead birds will also be underrepresented in this effort. Therefore, the real number of species that lack sufficient pathobiology data is likely larger than demonstrated here. This is partly because challenge studies with HPAI require BSL-3 laboratories with specialized spaces for avian species, which are limited and often have overriding study priorities. Additionally, it can be difficult to obtain and care for wild birds or wild bird eggs needed for conducting these experiments. Working to close this large data gap is a notable potential avenue for future research, even for species that are not suffering from large-scale mass mortality events and that do not have recognized roles in AIV spread.

## Supporting information

Review Database

Review Database Citaitons

## Acknowledgements

This work was funded by the U.S. Geological Survey, Ecosystem Mission Area. All data reported in this manuscript are publicly available [52]. Any use of trade, firm, or product names is for descriptive purposes only and does not imply endorsement by the U.S. Government.

