## Supplemental Tables and Figures for "A systematic review of laboratory investigations into the pathogenesis of avian influenza viruses in wild avifauna of North America"

Supplemental – Figures and Tables

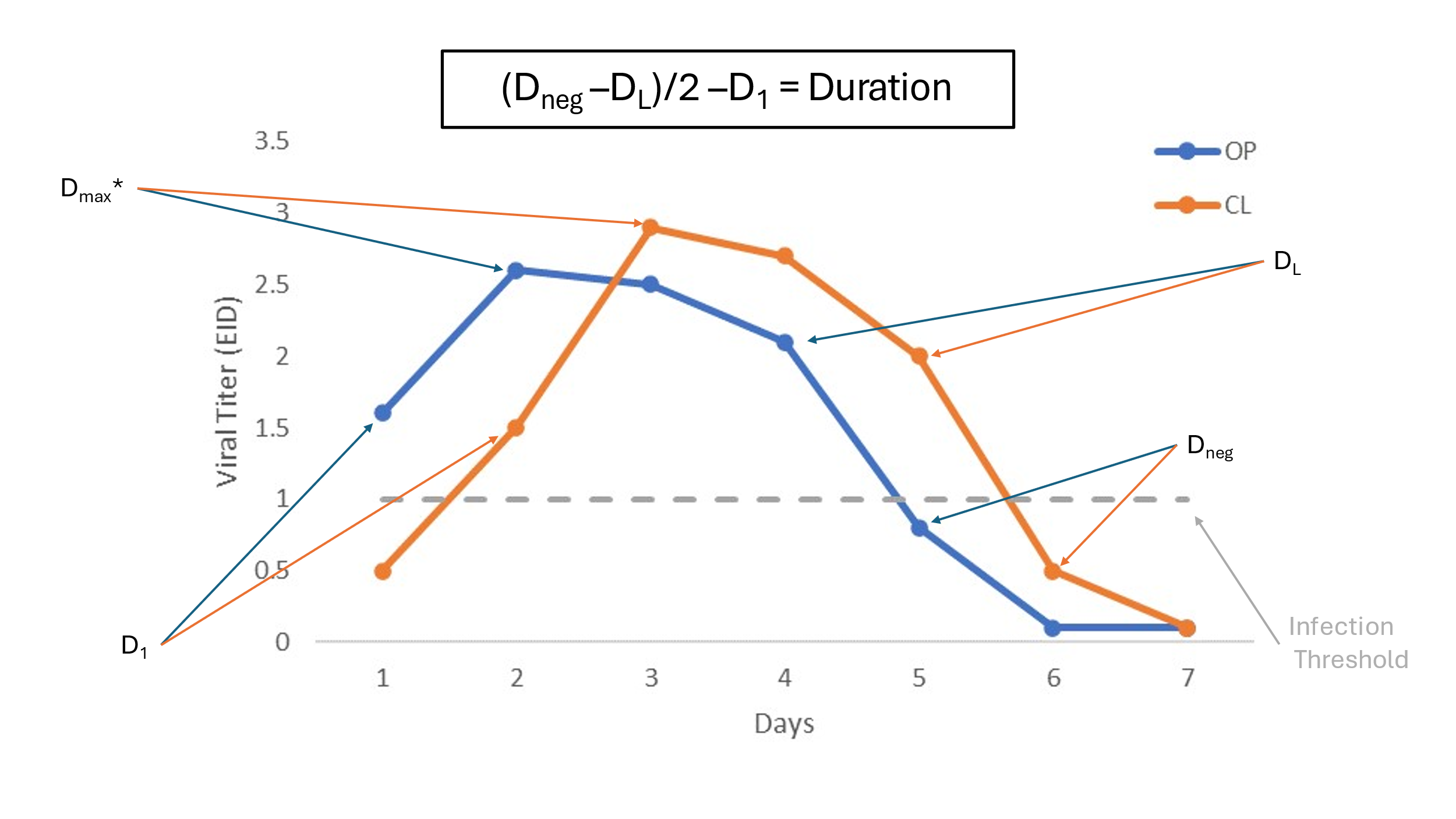

Figure S1. Example plot showing how we calculated duration of shedding if it was not provided within a publication. We identify specific time points describing the first day shedding was observed above a threshold value (D_1_), the greatest mean value on any given day (D_Max_), the last day shedding was observed above a given threshold (D_L_) and the first day animals tested consistently negative following infection (D_Neg_). In this plot viral titers are shown measured in EID (egg infectious dose) and were collected via oropharyngeal (OP) and cloacal swabs (CL).

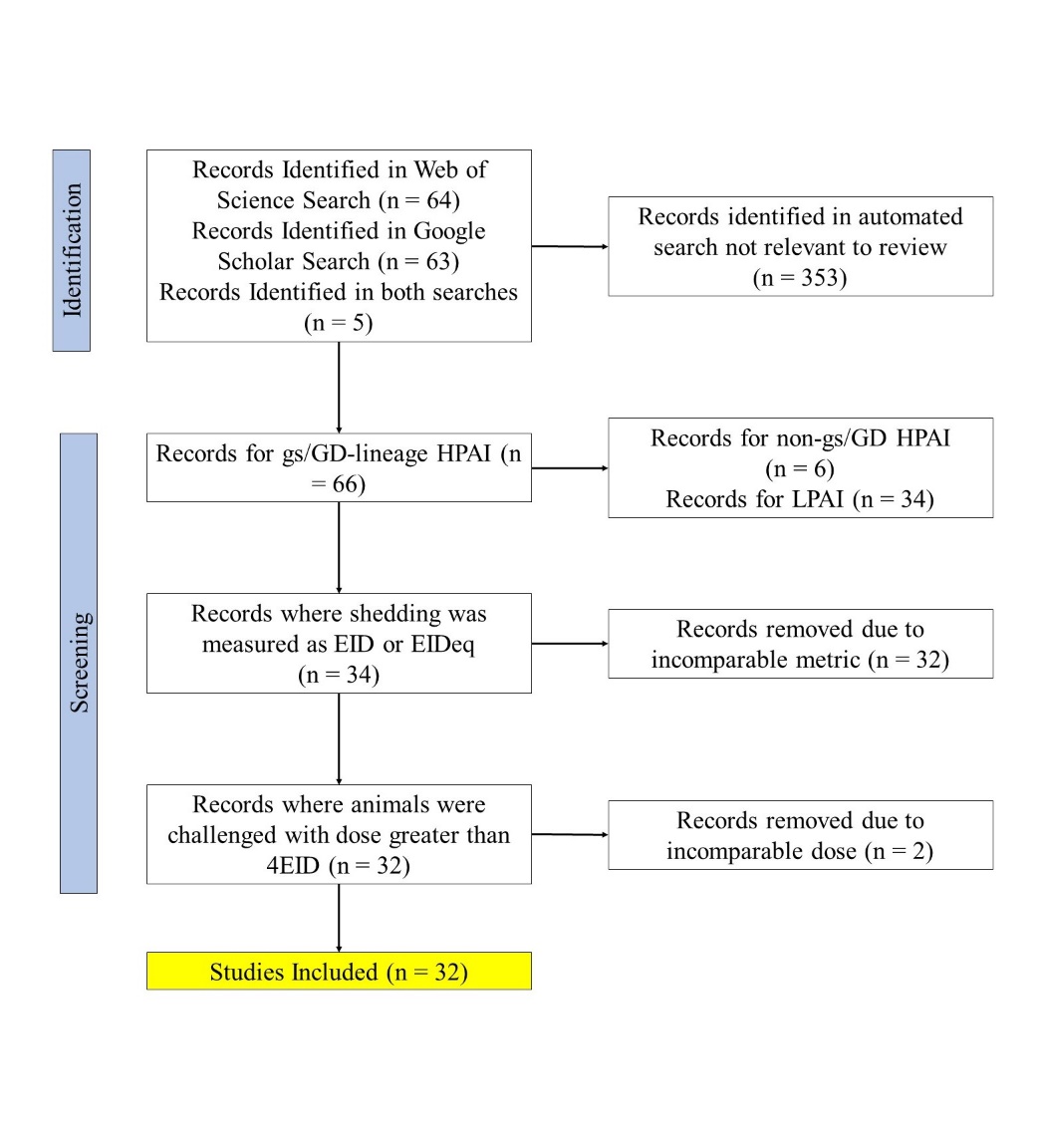

Figure S2. Flow chart showing the number of studies identified and screened through the course of the two literature reviews.

Table S1. Comparison of the number of studies and species per family identified in a literature review of avian influenza virus (AIV) challenge studies to the number of reported infections to the USDA-APHIS avian influenza monitoring database as of Nov 16, 2023 (USDA 2023). Viruses used include both highly pathogenic (HPAI) and low pathogenic avian influenza (LPAI).

|  |  | Studies | | | | Species | | |
| --- | --- | --- | --- | --- | --- | --- | --- | --- |
| Family | USDA Reports | LPAI | HPAI (Gs/GD) | HPAI (other) | LPAI | | HPAI (Gs/GD) | HPAI (other) |
| Accipitridae | 727 | 0 | 0 | 0 | 0 | | 0 | 0 |
| Alcidae | 20 | 0 | 0 | 0 | 0 | | 0 | 0 |
| Anatidae | 4667 | 37 | 46 | 8 | 9 | | 15 | 4 |
| Ardeidae | 22 | 0 | 1 | 0 | 0 | | 1 | 0 |
| Cathartidae | 691 | 0 | 0 | 0 | 0 | | 0 | 0 |
| Charadriidae | 4 | 0 | 0 | 0 | 0 | | 0 | 0 |
| Ciconiidae | 2 | 0 | 0 | 0 | 0 | | 0 | 0 |
| Columbidae | 0 | 2 | 5 | 1 | 1 | | 1 | 1 |
| Corvidae | 118 | 0 | 0 | 0 | 0 | | 0 | 0 |
| Falconidae | 46 | 0 | 2 | 0 | 0 | | 1 | 0 |
| Gaviidae | 9 | 0 | 0 | 0 | 0 | | 0 | 0 |
| Gruidae | 10 | 0 | 0 | 0 | 0 | | 0 | 0 |
| Hirundinidae | 1 | 0 | 0 | 0 | 0 | | 0 | 0 |
| Icteridae | 7 | 0 | 0 | 0 | 0 | | 0 | 0 |
| Laridae | 184 | 4 | 6 | 1 | 2 | | 3 | 1 |
| Pandionidae | 2 | 0 | 0 | 0 | 0 | | 0 | 0 |
| Passerellidae | 1 | 0 | 0 | 0 | 0 | | 0 | 0 |
| Passeridae | 1 | 3 | 5 | 0 | 1 | | 1 | 0 |
| Pelecanidae | 119 | 0 | 0 | 0 | 0 | | 0 | 0 |
| Phalacrocoracidae | 39 | 0 | 0 | 0 | 0 | | 0 | 0 |
| Phasianidae | 22 | 0 | 2 | 0 | 0 | | 1 | 0 |
| Podicipedidae | 18 | 0 | 0 | 0 | 0 | | 0 | 0 |
| Procellariidae | 4 | 0 | 0 | 0 | 0 | | 0 | 0 |
| Rallidae | 2 | 0 | 0 | 0 | 0 | | 0 | 0 |
| Scolopacidae | 28 | 2 | 3 | 0 | 1 | | 3 | 0 |
| Stercorariidae | 2 | 0 | 0 | 0 | 0 | | 0 | 0 |
| Strigidae | 248 | 0 | 0 | 0 | 0 | | 0 | 0 |
| Sulidae | 3 | 0 | 0 | 0 | 0 | | 0 | 0 |
| Threskiornithidae | 3 | 1 | 0 | 0 | 1 | | 0 | 0 |
| Turdidae | 1 | 0 | 1 | 0 | 0 | | 1 | 0 |
| Tytonidae | 3 | 0 | 0 | 0 | 0 | | 0 | 0 |

Table S2. Comparison of the number of studies by species identified in a literature review of avian influenza virus (AIV) challenge studies to the number of reported infections to the USDA-APHIS avian influenza monitoring database as of Nov 16, 2023 (USDA 2023).

|  |  |  |  | Studies | | |
| --- | --- | --- | --- | --- | --- | --- |
| Family | Scientific | Common | USDA Reports | LPAI | HPAI (gs/GD) | HPAI (other) |
| Accipitridae | *Accipiter cooperii* | Cooper's Hawk | 18 | 0 | 0 | 0 |
| Accipitridae | *Accipiter striatus* | Sharp-shinned Hawk | 6 | 0 | 0 | 0 |
| Accipitridae | *Aquila chrysaetos* | Golden Eagle | 8 | 0 | 0 | 0 |
| Accipitridae | *Buteo jamaicensis* | Red-tailed Hawk | 238 | 0 | 0 | 0 |
| Accipitridae | *Buteo lagopus* | Rough-legged Hawk | 6 | 0 | 0 | 0 |
| Accipitridae | *Buteo lineatus* | Red-shouldered Hawk | 9 | 0 | 0 | 0 |
| Accipitridae | *Buteo platypterus* | Broad-winged Hawk | 1 | 0 | 0 | 0 |
| Accipitridae | *Buteo regalis* | Ferruginous Hawk | 2 | 0 | 0 | 0 |
| Accipitridae | *Buteo swainsoni* | Swainson's Hawk | 8 | 0 | 0 | 0 |
| Accipitridae | *Circus hudsonius* | Northern Harrier | 2 | 0 | 0 | 0 |
| Accipitridae | *Haliaeetus leucocephalus* | Bald Eagle | 404 | 0 | 0 | 0 |
| Accipitridae | *Parabuteo unicinctus* | Harris's Hawk | 1 | 0 | 0 | 0 |
| Alcidae | *Uria aalge* | Common Murre | 18 | 0 | 0 | 0 |
| Anatidae | *Aix sponsa* | Wood Duck | 430 | 2 | 0 | 3 |
| Anatidae | *Anas acuta* | Northern Pintail | 104 | 0 | 0 | 4 |
| Anatidae | *Anas crecca* | American Green-winged Teal | 447 | 0 | 0 | 0 |
| Anatidae | *Anas fulvigula* | Mottled Duck | 1 | 0 | 0 | 0 |
| Anatidae | *Anas platyrhynchos* | Mallard | 927 | 28 | 5 | 29 |
| Anatidae | *Anas rubripes* | American Black Duck | 44 | 0 | 0 | 1 |
| Anatidae | *Anser albifrons* | Greater White-fronted Goose | 15 | 0 | 0 | 0 |
| Anatidae | *Anser caerulescens* | Snow Goose | 514 | 1 | 0 | 0 |
| Anatidae | *Anser rossii* | Ross's Goose | 102 | 0 | 0 | 0 |
| Anatidae | *Aythya affinis* | Lesser Scaup | 32 | 0 | 1 | 1 |
| Anatidae | *Aythya americana* | Redhead | 17 | 1 | 0 | 1 |
| Anatidae | *Aythya collaris* | Ring-necked Duck | 15 | 0 | 0 | 0 |
| Anatidae | *Aythya marila* | Greater Scaup | 1 | 0 | 0 | 0 |
| Anatidae | *Aythya valisineria* | Canvasback | 2 | 0 | 0 | 0 |
| Anatidae | *Branta bernicla* | Brant | 7 | 0 | 0 | 0 |
| Anatidae | *Branta canadensis* | Canada Goose | 577 | 3 | 0 | 5 |
| Anatidae | *Branta hutchinsii* | Cackling Goose | 32 | 0 | 0 | 1 |
| Anatidae | *Bucephala albeola* | Bufflehead | 9 | 0 | 0 | 0 |
| Anatidae | *Bucephala clangula* | Common Goldeneye | 6 | 0 | 0 | 0 |
| Anatidae | *Cairina moschata* | Muscovy Duck | 32 | 2 | 2 | 8 |
| Anatidae | *Cygnus buccinator* | Trumpeter Swan | 25 | 0 | 0 | 1 |
| Anatidae | *Cygnus columbianus* | Tundra swan | 11 | 0 | 0 | 0 |
| Anatidae | *Cygnus olor* | Mute Swan | 23 | 0 | 0 | 2 |
| Anatidae | *Dendrocygna bicolor* | Fulvous Whistling-Duck | 1 | 0 | 0 | 0 |
| Anatidae | *Lophodytes cucullatus* | Hooded Merganser | 23 | 0 | 0 | 0 |
| Anatidae | *Mareca americana* | American Wigeon | 278 | 0 | 0 | 0 |
| Anatidae | *Mareca strepera* | Gadwall | 226 | 0 | 0 | 1 |
| Anatidae | *Melanitta deglandi* | White-winged Scoter | 1 | 0 | 0 | 0 |
| Anatidae | *Melanitta perspicillata* | Surf Scoter | 0 | 0 | 0 | 1 |
| Anatidae | *Mergus merganser* | Common Merganser | 2 | 0 | 0 | 0 |
| Anatidae | *Oxyura jamaicensis* | Ruddy Duck | 6 | 0 | 0 | 1 |
| Anatidae | *Somateria mollissima* | Common Eider | 28 | 1 | 0 | 0 |
| Anatidae | *Spatula clypeata* | Northern Shoveler | 93 | 0 | 0 | 0 |
| Anatidae | *Spatula cyanoptera* | Cinnamon Teal | 20 | 1 | 1 | 0 |
| Anatidae | *Spatula discors* | Blue-winged Teal | 504 | 2 | 0 | 1 |
| Ardeidae | *Ardea alba* | Great Egret | 3 | 0 | 0 | 0 |
| Ardeidae | *Ardea herodias* | Great Blue Heron | 9 | 0 | 0 | 0 |
| Ardeidae | *Butorides virescens* | Green Heron | 1 | 0 | 0 | 0 |
| Ardeidae | *Egretta thula* | Snowy Egret | 3 | 0 | 0 | 0 |
| Ardeidae | *Nycticorax nycticorax* | Black-crowned Night Heron | 5 | 0 | 0 | 1 |
| Cathartidae | *Cathartes aura* | Turkey Vulture | 117 | 0 | 0 | 0 |
| Cathartidae | *Coragyps atratus* | Black Vulture | 550 | 0 | 0 | 0 |
| Cathartidae | *Gymnogyps californianus* | California Condor | 22 | 0 | 0 | 0 |
| Charadriidae | *Charadrius nivosus* | Snowy Plover | 4 | 0 | 0 | 0 |
| Ciconiidae | *Mycteria americana* | American Wood Stork | 2 | 0 | 0 | 0 |
| Columbidae | *Columba livia* | Rock Pigeon | 0 | 2 | 1 | 5 |
| Corvidae | *Corvus brachyrhynchos* | American Crow | 64 | 0 | 0 | 0 |
| Corvidae | *Corvus corax* | Common Raven | 36 | 0 | 0 | 0 |
| Corvidae | *Corvus ossifragus* | Fish Crow | 5 | 0 | 0 | 0 |
| Corvidae | *Pica hudsonia* | Black-billed Magpie | 3 | 0 | 0 | 0 |
| Falconidae | *Caracara cheriway* | Crested Caracara | 1 | 0 | 0 | 0 |
| Falconidae | *Falco columbarius* | Merlin | 1 | 0 | 0 | 0 |
| Falconidae | *Falco mexicanus* | Prairie Falcon | 1 | 0 | 0 | 0 |
| Falconidae | *Falco peregrinus* | Peregrine Falcon | 42 | 0 | 0 | 0 |
| Falconidae | *Falco sparverius* | American Kestrel | 1 | 0 | 0 | 2 |
| Gaviidae | *Gavia immer* | Common Loon | 8 | 0 | 0 | 0 |
| Gaviidae | *Gavia pacifica* | Pacific Loon | 1 | 0 | 0 | 0 |
| Gruidae | *Antigone canadensis* | Sandhill Crane | 10 | 0 | 0 | 0 |
| Hirundinidae | *Tachycineta bicolor* | Tree Swallow | 1 | 0 | 0 | 0 |
| Icteridae | *Agelaius phoeniceus* | Red-winged Blackbird | 1 | 0 | 0 | 0 |
| Icteridae | *Quiscalus major* | Boat-tailed Grackle | 1 | 0 | 0 | 0 |
| Icteridae | *Quiscalus mexicanus* | Great-tailed Grackle | 3 | 0 | 0 | 0 |
| Icteridae | *Quiscalus quiscula* | Common Grackle | 2 | 0 | 0 | 0 |
| Laridae | *Chroicocephalus philadelphia* | Bonaparte's Gull | 1 | 0 | 0 | 0 |
| Laridae | *Chroicocephalus ridibundus* | Black-headed Gull | 0 | 0 | 0 | 1 |
| Laridae | *Hydroprogne caspia* | Caspian Tern | 26 | 0 | 0 | 0 |
| Laridae | *Larus argentatus* | Herring Gull | 36 | 0 | 0 | 2 |
| Laridae | *Larus brachyrhynchus* | Short-billed Gull | 1 | 0 | 0 | 0 |
| Laridae | *Larus californicus* | California Gull | 2 | 0 | 0 | 0 |
| Laridae | *Larus delawarensis* | Ring-billed Gull | 6 | 1 | 0 | 0 |
| Laridae | *Larus glaucescens* | Glaucous-winged Gull | 3 | 0 | 0 | 0 |
| Laridae | *Larus glaucoides thayeri* | Thayer's Gull | 1 | 0 | 0 | 0 |
| Laridae | *Larus hyperboreus* | Glaucous Gull | 14 | 0 | 0 | 0 |
| Laridae | *Larus marinus* | Great Black-backed Gull | 29 | 0 | 0 | 0 |
| Laridae | *Larus occidentalis* | Western Gull | 12 | 0 | 0 | 0 |
| Laridae | *Leucophaeus atricilla* | Laughing Gull | 1 | 3 | 1 | 3 |
| Laridae | *Rissa tridactyla* | Black-legged Kittiwake | 2 | 0 | 0 | 0 |
| Laridae | *Rynchops niger* | Black Skimmer | 4 | 0 | 0 | 0 |
| Laridae | *Sterna forsteri* | Forster's Tern | 1 | 0 | 0 | 0 |
| Laridae | *Sterna hirundo* | Common Tern | 11 | 0 | 0 | 0 |
| Laridae | *Sterna paradisaea* | Arctic Tern | 2 | 0 | 0 | 0 |
| Laridae | *Thalasseus maximus* | Royal Tern | 3 | 0 | 0 | 0 |
| Laridae | *Thalasseus sandvicensis* | Sandwich Tern | 1 | 0 | 0 | 0 |
| Laridae | *Xema sabini* | Sabine's Gull | 3 | 0 | 0 | 0 |
| Pandionidae | *Pandion haliaetus* | Osprey | 2 | 0 | 0 | 0 |
| Passerellidae | *Junco hyemalis* | Dark-eyed junco | 1 | 0 | 0 | 0 |
| Passeridae | *Passer domesticus* | House Sparrow | 1 | 3 | 0 | 5 |
| Pelecanidae | *Pelecanus erythrorhynchos* | American White Pelican | 97 | 0 | 0 | 0 |
| Pelecanidae | *Pelecanus occidentalis* | Brown Pelican | 13 | 0 | 0 | 0 |
| Phalacrocoracidae | *Nannopterum auritum* | Double-crested Cormorant | 27 | 0 | 0 | 0 |
| Phalacrocoracidae | *Nannopterum brasilianum* | Neotropic Cormorant | 3 | 0 | 0 | 0 |
| Phalacrocoracidae | *Urile penicillatus* | Brandt's Cormorant | 1 | 0 | 0 | 0 |
| Phasianidae | *Bonasa umbellus* | Ruffed Grouse | 1 | 0 | 0 | 0 |
| Phasianidae | *Centrocercus urophasianus* | Greater Sage-grouse | 1 | 0 | 0 | 0 |
| Phasianidae | *Meleagris gallopavo* | Wild turkey | 17 | 0 | 0 | 0 |
| Phasianidae | *Phasianus colchicus* | Ringed-neck pheasant | 0 | 0 | 0 | 2 |
| Podicipedidae | *Aechmophorus occidentalis* | Western Grebe | 1 | 0 | 0 | 0 |
| Podicipedidae | *Podiceps auritus* | Horned Grebe | 2 | 0 | 0 | 0 |
| Podicipedidae | *Podiceps grisegena* | Red-necked Grebe | 1 | 0 | 0 | 0 |
| Podicipedidae | *Podiceps nigricollis* | Eared Grebe | 13 | 0 | 0 | 0 |
| Podicipedidae | *Podilymbus podiceps* | Pied-billed Grebe | 1 | 0 | 0 | 0 |
| Procellariidae | *Ardenna tenuirostris* | Short-tailed Shearwater | 3 | 0 | 0 | 0 |
| Procellariidae | *Fulmarus glacialis* | Northern Fulmar | 1 | 0 | 0 | 0 |
| Rallidae | *Fulica americana* | American Coot | 2 | 0 | 0 | 0 |
| Scolopacidae | *Arenaria interpres* | Ruddy Turnstone | 1 | 2 | 0 | 1 |
| Scolopacidae | *Arenaria melanocephala* | Black Turnstone | 2 | 0 | 0 | 0 |
| Scolopacidae | *Calidris alba* | Sanderling | 20 | 0 | 0 | 0 |
| Scolopacidae | *Calidris alpina* | Dunlin | 3 | 0 | 0 | 1 |
| Scolopacidae | *Calidris canutus* | Red Knot | 0 | 0 | 0 | 1 |
| Scolopacidae | *Phalaropus lobatus* | Red-necked Phalarope | 1 | 0 | 0 | 0 |
| Scolopacidae | *Tringa semipalmata* | Willet | 1 | 0 | 0 | 0 |
| Stercorariidae | *Stercorarius parasiticus* | Parasitic Jaeger | 2 | 0 | 0 | 0 |
| Strigidae | *Asio flammeus* | Short-eared Owl | 1 | 0 | 0 | 0 |
| Strigidae | *Asio otus* | Long-eared Owl | 1 | 0 | 0 | 0 |
| Strigidae | *Bubo scandiacus* | Snowy Owl | 15 | 0 | 0 | 0 |
| Strigidae | *Bubo virginianus* | Great Horned Owl | 204 | 0 | 0 | 0 |
| Strigidae | *Megascops asio* | Eastern Screech-Owl | 4 | 0 | 0 | 0 |
| Strigidae | *Megascops kennicottii* | Western Screech-Owl | 2 | 0 | 0 | 0 |
| Strigidae | *Strix varia* | Barred Owl | 10 | 0 | 0 | 0 |
| Sulidae | *Morus bassanus* | Northern Gannet | 2 | 0 | 0 | 0 |
| Threskiornithidae | *Eudocimus albus* | White Ibis | 1 | 1 | 0 | 0 |
| Threskiornithidae | *Platalea ajaja* | Roseate Spoonbill | 1 | 0 | 0 | 0 |
| Threskiornithidae | *Plegadis falcinellus* | Glossy Ibis | 1 | 0 | 0 | 0 |
| Turdidae | *Turdus migratorius* | American Robin | 1 | 0 | 0 | 1 |
| Tytonidae | *Tyto alba* | Barn Owl | 3 | 0 | 0 | 0 |

Table S3. Summary of infection rates for avian species directly inoculated with greater than 4 log_10_ EID_50_/mL of Gs/GD HPAI virus, reported by host species. We report the number of studies identified (Studies), total number of individuals challenged (Sampled), the number removed according to study design (Culled), the total number and percentage of challenged individuals infected (Infection) and the total number of individuals that died after accounting for the number culled (Mortality).

|  |  |  |  |  | Infection | |  | Mortality | |
| --- | --- | --- | --- | --- | --- | --- | --- | --- | --- |
| Family | Scientific | Common | Studies | Sampled | Total | Rate | Culled | Total | Rate |
| Anatidae | *Aix sponsa* | Wood Duck | 3 | 41 | 41 | 1.00 | 0 | 30 | 0.73 |
| Anatidae | *Anas acuta* | Northern Pintail | 4 | 30 | 24 | 0.80 | 6 | 0 | 0.00 |
| Anatidae | *Anas platyrhynchos* | Mallard | 24 | 702 | 520 | 0.74 | 161 | 166 | 0.31 |
| Anatidae | *Anas rubripes* | American Black Duck | 1 | 16 | 16 | 1.00 | 0 | 0 | 0.00 |
| Anatidae | *Aythya affinis* | Lesser Scaup | 1 | 18 | 18 | 1.00 | 0 | 0 | 0.00 |
| Anatidae | *Aythya americana* | Redhead | 1 | 6 | 5 | 0.83 | 0 | 0 | 0.00 |
| Anatidae | *Branta canadensis* | Canada Goose | 1 | 20 | 20 | 1.00 | 0 | 20 | 1.00 |
| Anatidae | *Branta hutchinsii* | Cackling Goose | 1 | 4 | 3 | 0.75 | 0 | 3 | 0.75 |
| Anatidae | *Cairina moschata* | Muscovy Duck | 8 | 216 | 204 | 0.94 | 57 | 115 | 0.72 |
| Anatidae | *Cygnus buccinator* | Trumpeter Swan | 1 | 5 | 5 | 1.00 | 0 | 5 | 1.00 |
| Anatidae | *Cygnus olor* | Mute Swan | 2 | 7 | 7 | 1.00 | 0 | 7 | 1.00 |
| Anatidae | *Melanitta perspicillata* | Surf Scoter | 1 | 9 | 7 | 0.78 | 0 | 2 | 0.22 |
| Anatidae | *Oxyura jamaicensis* | Ruddy Duck | 1 | 20 | 19 | 0.95 | 0 | 4 | 0.20 |
| Anatidae | *Spatula discors* | Blue-winged Teal | 1 | 6 | 5 | 0.83 | 0 | 0 | 0.00 |
| Ardeidae | *Nycticorax nycticorax* | Black-crowned Night Heron | 1 | 3 | 1 | 0.33 | 0 | 1 | 0.33 |
| Columbidae | *Columba livia* | Rock Pigeon | 3 | 67 | 20 | 0.30 | 38 | 2 | 0.07 |
| Falconidae | *Falco sparverius* | American Kestrel | 2 | 8 | 8 | 1.00 | 0 | 8 | 1.00 |
| Laridae | *Larus argentatus* | Herring Gull | 2 | 42 | 42 | 1.00 | 0 | 28 | 0.67 |
| Laridae | *Leucophaeus atricilla* | Laughing Gull | 3 | 22 | 14 | 0.64 | 8 | 4 | 0.29 |
| Passeridae | *Passer domesticus* | House Sparrow | 4 | 36 | 28 | 0.78 | 0 | 21 | 0.58 |
| Phasianidae | *Phasianus colchicus* | Ringed-neck pheasant | 2 | 20 | 20 | 1.00 | 0 | 20 | 1.00 |
| Scolopacidae | *Arenaria interpres* | Ruddy Turnstone | 1 | 17 | 14 | 0.82 | 0 | 0 | 0.00 |
| Scolopacidae | *Calidris alpina* | Dunlin | 1 | 6 | 6 | 1.00 | 0 | 6 | 1.00 |
| Turdidae | *Turdus migratorius* | American Robin | 1 | 24 | 22 | 0.92 | 0 | 0 | 0.00 |

Table S4. Summary of viral shedding for avian species directly inoculated with greater than 4 log_10_ EID_50_/mL of Gs/GD HPAI virus and reported as EID_50_/mL or an EID_50_/mL equivalent (eqEID). We report the number of studies identified, the number of unique species challenged, the duration in days and the viral shedding rates measured using virus isolation or RT-PCR methods. We report information separately for oropharyngeal (OP) and cloacal (CL) pathways.

|  |  |  |  | Viral Shedding (EID_50_/mL) | | | Duration (Days) | |
| --- | --- | --- | --- | --- | --- | --- | --- | --- |
| Family | Scientific | Common | Studies | Orpharyngeal | Cloacal | Oropharyngeal | | Cloacal |
| Anatidae | *Aix sponsa* | Wood Duck | 3 | 4.79 EID (4.43 - 5.14) | 3.31 EID  (2.8 - 3.8) | 7.67 Days  (6 - 10) | | 3.67 Day (3 - 5) |
| Anatidae | *Anas acuta* | Northern Pintail | 3 | 2.22 EID (1.1 - 3.92) | 1 EID | 2 Days  (0.5 - 4) | | 1 Days |
| Anatidae | *Anas platyrhynchos* | Mallard | 21 | 4.78 EID (1.8 - 7.7) | 3.52 EID  (0.9 - 6.8) | 6.42 Days  (1 - 13) | | 6.03 Day (0.5 - 13) |
| Anatidae | *Anas rubripes* | American Black Duck | 1 | 4.47 EID (4 - 5.1) | 2.45 EID  (2.3 - 2.6) | 6.25 Days  (3.5 - 9) | | 6.5 Days |
| Anatidae | *Aythya affinis* | Lesser Scaup | 1 | - | - | 2 Days | | 2 Days |
| Anatidae | *Aythya americana* | Redhead | 1 | 3.4 EID (2.8 - 4) | 1.2 EID | 4 Days | | 1 Days |
| Anatidae | *Branta hutchinsii* | Cackling Goose | 1 | 5.25 EID | 3.05 EID | 5 Days | | 3 Days |
| Anatidae | *Cairina moschata* | Muscovy Duck | 8 | 5.31 EID (2.5 - 6.8) | 3.1 EID  (2.3 - 4.6) | 4.33 Days  (3 - 8) | | 4.38 Day (3 - 7) |
| Anatidae | *Cygnus buccinator* | Trumpeter Swan | 1 | 6.14 EID | 3.18 EID | 5 Days | | 4 Days |
| Anatidae | *Cygnus olor* | Mute Swan | 1 | 5.58 EID | 4.46 EID | 5 Days | | 4 Days |
| Anatidae | *Melanitta perspicillata* | Surf Scoter | 1 | 6.1 EID | 2.9 EID | 8 Days | | 8 Days |
| Anatidae | *Oxyura jamaicensis* | Ruddy Duck | 1 | - | - | 2 Days | | 2 Days |
| Anatidae | *Spatula discors* | Blue-winged Teal | 1 | 2.9 EID (2 - 3.8) | 1 EID | 2 Days | | 1 Days |
| Ardeidae | *Nycticorax nycticorax* | Black-crowned Night Heron | 1 | 4.3 EID | 2.5 EID | 3.5 Days | | 3.5 Days |
| Columbidae | *Columba livia* | Rock Pigeon | 2 | 2.15 EID | - | - | | - |
| Laridae | *Larus argentatus* | Herring Gull | 2 | 3.82 EID (3.74 - 3.89) | 2.01 EID  (1.96 - 2.07) | 5.27 Days  (4.3 - 6.5) | | 2.93 Day (2.3 - 3.5) |
| Laridae | *Leucophaeus atricilla* | Laughing Gull | 3 | 3.48 EID (1.24 - 5) | 1.86 EID  (0.97 - 2.6) | 9 Days  (8 - 10) | | 6.5 Day (6 - 7) |
| Passeridae | *Passer domesticus* | House Sparrow | 4 | 3.28 EID (2.6 - 4.7) | 2.7 EID  (1.3 - 4.1) | 4.8 Days  (4 - 6) | | 4.78 Day (3.95 - 6) |
| Phasianidae | *Phasianus colchicus* | Ringed-neck pheasant | 2 | - | - | 6 Days  (4 - 8) | | 6 Day (4 - 8) |
| Scolopacidae | *Calidris alpina* | Dunlin | 1 | 3.54 EID | 3.54 EID | 4 Days | | 4 Days |
| Turdidae | *Turdus migratorius* | American Robin | 1 | - | - | 3.83 Days  (3.5 - 4.5) | | - |

Tables S5. Summary of viral shedding for avian species challenged with Gs/GD highly pathogenic avian influenza (HPAI) virus and reported as egg infectious dose (EID_50_/mL) or an EID_50_/mL equivalent (eqEID). Information is reported by host species and virus lineage. We report the number of studies identified, the number of unique species challenged, the duration in days and the viral shedding rates measured using virus isolation or reverse transcriptase polymerase chain reaction (RT-PCR) methods. We report this information separately for the oropharyngeal (OP) and cloacal (CL) pathways.

|  |  |  |  | Quantity (EID50/mL) | Duration (Days) |  |  |
| --- | --- | --- | --- | --- | --- | --- | --- |
| Family | Scientific | Common | Studies | OP | CL | OP | CL |
| Anatidae | *Aix sponsa* | Wood Duck | 3 | 4.76 EID (4.38 - 5.14) | 2.99 EID  (1.96 - 3.8) | 9 Days  (6 - 10) | 3.29 Day (3 - 5) |
| Anatidae | *Anas acuta* | Northern Pintail | 4 | 2.89 EID (1.1 - 4.6) | 1 EID | 2.44 Days  (0.5 - 5) | 1.67 Day (1 - 3) |
| Anatidae | *Anas platyrhynchos* | Mallard | 29 | 4.7 EID (1.5 - 7.7) | 3.41 EID  (0.7 - 6.8) | 5.83 Days  (0.5 - 13) | 5.74 Day (0.5 - 14) |
| Anatidae | *Anas rubripes* | American Black Duck | 1 | 3.97 EID (1.3 - 5.1) | 2.45 EID  (2.3 - 2.6) | 5.83 Days  (1 - 9) | 6.5 Days |
| Anatidae | *Aythya affinis* | Lesser Scaup | 1 | - | - | 2 Days | 2 Days |
| Anatidae | *Aythya americana* | Redhead | 1 | 3.4 EID (2.8 - 4) | 1.2 EID | 4 Days | 1 Days |
| Anatidae | *Branta canadensis* | Canada Goose | 5 | 5.88 EID (3.6 - 7.4) | 5.75 EID  (3.8 - 7.3) | 3.17 Days  (2 - 4) | 2.58 Day (1.5 - 4) |
| Anatidae | *Branta hutchinsii* | Cackling Goose | 1 | 5.25 EID | 3.05 EID | 5 Days | 3 Days |
| Anatidae | *Cairina moschata* | Muscovy Duck | 8 | 4.99 EID (2.42 - 6.8) | 3.11 EID  (1.92 - 4.6) | 4.07 Days  (3 - 8) | 4.46 Day (3 - 7) |
| Anatidae | *Cygnus buccinator* | Trumpeter Swan | 1 | 6.14 EID | 3.18 EID | 5 Days | 4 Days |
| Anatidae | *Cygnus olor* | Mute Swan | 2 | 5.58 EID | 4.46 EID | 5 Days | 4 Days |
| Anatidae | *Mareca strepera* | Gadwall | 1 | - | - | 1 Days | - |
| Anatidae | *Melanitta perspicillata* | Surf Scoter | 1 | 6.1 EID | 2.9 EID | 8 Days | 8 Days |
| Anatidae | *Oxyura jamaicensis* | Ruddy Duck | 1 | - | - | 2 Days | 2 Days |
| Anatidae | *Spatula discors* | Blue-winged Teal | 1 | 2.9 EID (2 - 3.8) | 1 EID | 2 Days | 1 Days |
| Ardeidae | *Nycticorax nycticorax* | Black-crowned Night Heron | 1 | 4.3 EID | 2.5 EID | 3.5 Days | 3.5 Days |
| Columbidae | *Columba livia* | Rock Pigeon | 5 | 2.15 EID | - | 1 Days | 1 Days |
| Falconidae | *Falco sparverius* | American Kestrel | 2 | 5.4 EID | 3.5 EID | 5.2 Days  (3 - 6) | 3 Day (2 - 4) |
| Laridae | *Chroicocephalus ridibundus* | Black-headed Gull | 1 | - | - | 12 Days | 11 Days |
| Laridae | *Larus argentatus* | Herring Gull | 2 | 4.03 EID (3.74 - 4.47) | 2.14 EID  (1.96 - 2.38) | 6.13 Days  (3.5 - 13) | 4.63 Day (2.3 - 13) |
| Laridae | *Leucophaeus atricilla* | Laughing Gull | 3 | 3.48 EID (1.24 - 5) | 1.86 EID  (0.97 - 2.6) | 9 Days  (8 - 10) | 6.5 Day (6 - 7) |
| Passeridae | *Passer domesticus* | House Sparrow | 5 | 2.96 EID (2.3 - 4.7) | 2.7 EID  (1.3 - 4.1) | 4.48 Days  (3 - 6) | 4.77 Day (3.95 - 6) |
| Phasianidae | *Phasianus colchicus* | Ringed-neck pheasant | 2 | - | - | 5.12 Days  (2 - 8) | 4.62 Day (3 - 8) |
| Scolopacidae | *Arenaria interpres* | Ruddy Turnstone | 1 | - | - | 6.5 Days  (5.5 - 8.5) | - |
| Scolopacidae | *Calidris alpina* | Dunlin | 1 | 3.24 EID (3.06 - 3.54) | 3.11 EID  (2.67 - 3.54) | 4.26 Days  (4 - 4.67) | 3.9 Day (3 - 4.71) |
| Scolopacidae | *Calidris canutus* | Red Knot | 1 | - | - | 3.5 Days | - |
| Turdidae | *Turdus migratorius* | American Robin | 1 | - | - | 3.83 Days  (3.5 - 4.5) | - |

This work was funded by the U.S. Geological Survey, Ecosystem Mission Area. All data reported in this manuscript are publicly available. Any use of trade, firm, or product names is for descriptive purposes only and does not imply endorsement by the U.S. Government.
