## Supplementary material for "A systematic review of laboratory investigations into the pathogenesis of avian influenza viruses in wild avifauna of North America": Review Database Citaitons

Supplemental 1 – Citations

Aamir, Uzma B., Khalid Naeem, Zaheer Ahmed, Caroline A. Obert, John Franks, Scott Krauss, Patrick Seiler, and Robert G. Webster. “Zoonotic Potential of Highly Pathogenic Avian H7N3 Influenza Viruses from Pakistan.” *Virology* 390, no. 2 (August 1, 2009): 212–20. <https://doi.org/10.1016/j.virol.2009.05.008>.

Ahrens, Ann Kathrin, Hans-Christoph Selinka, Thomas C. Mettenleiter, Martin Beer, and Timm C. Harder. “Exploring Surface Water as a Transmission Medium of Avian Influenza Viruses – Systematic Infection Studies in Mallards.” *Emerging Microbes & Infections* 11, no. 1 (December 31, 2022): 1250–61. <https://doi.org/10.1080/22221751.2022.2065937>.

Arsnoe, Dustin M., Hon S. Ip, and Jennifer C. Owen. “Influence of Body Condition on Influenza A Virus Infection in Mallard Ducks: Experimental Infection Data.” *PLOS ONE* 6, no. 8 (August 16, 2011): e22633. <https://doi.org/10.1371/journal.pone.0022633>.

Bahnson, Charlie S., Sonia M. Hernandez, Rebecca L. Poulson, Robert E. Cooper, Shannon E. Curry, Taylor J. Ellison, Henry C. Adams, Catharine N. Welch, and David E. Stallknecht. “EXPERIMENTAL INFECTIONS AND SEROLOGY INDICATE THAT AMERICAN WHITE IBIS (EUDOCIUMUS ALBUS) ARE COMPETENT RESERVOIRS FOR TYPE A INFLUENZA VIRUS.” *Journal of Wildlife Diseases* 56, no. 3 (July 1, 2020): 530–37. <https://doi.org/10.7589/2019-05-136>.

Bahnson, Charlie S., Rebecca L. Poulson, Laura P. Hollander, Jo A. Crum Bradley, and David E. Stallknecht. “SUSCEPTIBILITY OF LAUGHING GULLS (LEUCOPHAEUS ATRICILLA) AND MALLARDS (ANAS PLATYRHYNCHOS) TO RUDDY TURNSTONE (ARENARIA INTERPRES MORINELLA) ORIGIN TYPE A INFLUENZA VIRUSES.” *Journal of Wildlife Diseases* 56, no. 1 (January 2020): 167–74. <https://doi.org/10.7589/2019-03-065>.

Berhane, Y., M. Leith, C. Embury-Hyatt, J. Neufeld, S. Babiuk, T. Hisanaga, H. Kehler, K. Hooper-McGrevy, and J. Pasick. “Studying Possible Cross-Protection of Canada Geese Preexposed to North American Low Pathogenicity Avian Influenza Virus Strains (H3N8, H4N6, and H5N2) Against an H5N1 Highly Pathogenic Avian Influenza Challenge.” *Avian Diseases* 54, no. s1 (March 2010): 548–54. <https://doi.org/10.1637/8841-040309-Reg.1>.

Berhane, Yohannes, Carissa Embury-Hyatt, Marsha Leith, Helen Kehler, Matthew Suderman, and John Pasick. “PRE-EXPOSING CANADA GEESE (BRANTA CANADENSIS) TO A LOW-PATHOGENIC H1N1 AVIAN INFLUENZA VIRUS PROTECTS THEM AGAINST H5N1 HPAI VIRUS CHALLENGE.” *Journal of Wildlife Diseases* 50, no. 1 (January 2014): 84–97. <https://doi.org/10.7589/2012-09-237>.

Bertran, Kateri, Mary J. Pantin-Jackwood, Miria F. Criado, Dong-Hun Lee, Charles L. Balzli, Erica Spackman, David L. Suarez, and David E. Swayne. “Pathobiology and Innate Immune Responses of Gallinaceous Poultry to Clade 2.3.4.4A H5Nx Highly Pathogenic Avian Influenza Virus Infection.” *Veterinary Research* 50, no. 1 (November 1, 2019): 89. <https://doi.org/10.1186/s13567-019-0704-5>.

Bertran, Kateri, Elisa Pérez-Ramírez, Núria Busquets, Roser Dolz, Antonio Ramis, Ayub Darji, Francesc Xavier Abad, et al. “Pathogenesis and Transmissibility of Highly (H7N1) and Low (H7N9) Pathogenic Avian Influenza Virus Infection in Red-Legged Partridge (Alectoris Rufa).” *Veterinary Research* 42, no. 1 (February 7, 2011): 24. <https://doi.org/10.1186/1297-9716-42-24>.

Boon, Adrianus C. M., Matthew R. Sandbulte, Patrick Seiler, Richard J. Webby, Thaweesak Songserm, Yi Guan, and Robert G. Webster. “Role of Terrestrial Wild Birds in Ecology of Influenza A Virus (H5N1) - Volume 13, Number 11—November 2007 - Emerging Infectious Diseases Journal - CDC,” 2007. <https://doi.org/10.3201/eid1311.070114>.

Bosco-Lauth, Angela M., Nicole L. Marlenee, Airn E. Hartwig, Richard A. Bowen, and J. Jeffrey Root. “Shedding of Clade 2.3.4.4 H5N8 and H5N2 Highly Pathogenic Avian Influenza Viruses in Peridomestic Wild Birds in the U.S.” *Transboundary and Emerging Diseases* 66, no. 3 (2019): 1301–5. <https://doi.org/10.1111/tbed.13147>.

Bosco-Lauth, Angela, Anna Rodriguez, Rachel M. Maison, Stephanie M. Porter, and J. Jeffrey Root. “H7N9 Influenza A Virus Transmission in a Multispecies Barnyard Model.” *Virology* 582 (May 2023): 100–105. <https://doi.org/10.1016/j.virol.2023.04.002>.

Brand, Judith M. A. van den, Josanne H. Verhagen, Edwin J. B. Veldhuis Kroeze, Marco W. G. van de Bildt, Rogier Bodewes, Sander Herfst, Mathilde Richard, et al. “Wild Ducks Excrete Highly Pathogenic Avian Influenza Virus H5N8 (2014–2015) without Clinical or Pathological Evidence of Disease.” *Emerging Microbes & Infections* 7, no. 1 (December 1, 2018): 1–10. <https://doi.org/10.1038/s41426-018-0070-9>.

Bröjer, Caroline, Josef D. Järhult, Shaman Muradrasoli, Hanna Söderstrom, Björn Olsen, and Dolores Gavier-Widén. “PATHOBIOLOGY AND VIRUS SHEDDING OF LOW-PATHOGENIC AVIAN INFLUENZA VIRUS (A/H1N1) INFECTION IN MALLARDS EXPOSED TO OSELTAMIVIR.” *Journal of Wildlife Diseases* 49, no. 1 (January 2013): 103–13. <https://doi.org/10.7589/2011-11-335>.

Brown, Justin D., Roy D. Berghaus, Taiana P. Costa, Rebecca Poulson, Deborah L. Carter, Camille Lebarbenchon, and David E. Stallknecht. “INTESTINAL EXCRETION OF A WILD BIRD-ORIGIN H3N8 LOW PATHOGENIC AVIAN INFLUENZA VIRUS IN MALLARDS (ANAS PLATYRHYNCHOS).” *Journal of Wildlife Diseases* 48, no. 4 (October 2012): 991–98. <https://doi.org/10.7589/2011-09-280>.

Brown, Justin D., David E. Stallknecht, Joan R. Beck, David L. Suarez, and David E. Swayne. “Susceptibility of North American Ducks and Gulls to H5N1 Highly Pathogenic Avian Influenza Viruses.” *Emerging Infectious Diseases* 12, no. 11 (November 2006): 1663–70. <https://doi.org/10.3201/eid1211.060652>.

Brown, Justin D., David E. Stallknecht, Roy D. Berghaus, and David E. Swayne. “Infectious and Lethal Doses of H5N1 Highly Pathogenic Avian Influenza Virus for House Sparrows (Passer Domesticus) and Rock Pigeons (Columbia Livia).” *Journal of Veterinary Diagnostic Investigation* 21, no. 4 (July 1, 2009): 437–45. <https://doi.org/10.1177/104063870902100404>.

Brown, Justin D., David E. Stallknecht, and David E. Swayne. “Experimental Infection of Swans and Geese with Highly Pathogenic Avian Influenza Virus (H5N1) of Asian Lineage - Volume 14, Number 1—January 2008 - Emerging Infectious Diseases Journal - CDC,” 2008. <https://doi.org/10.3201/eid1401.070740>.

———. “Experimental Infections of Herring Gulls (Larus Argentatus) with H5N1 Highly Pathogenic Avian Influenza Viruses by Intranasal Inoculation of Virus and Ingestion of Virus-Infected Chicken Meat.” *Avian Pathology* 37, no. 4 (August 1, 2008): 393–97. <https://doi.org/10.1080/03079450802216595>.

Brown, Justin D., David E. Stallknecht, Steve Valeika, and David E. Swayne. “SUSCEPTIBILITY OF WOOD DUCKS TO H5N1 HIGHLY PATHOGENIC AVIAN INFLUENZA VIRUS.” *Journal of Wildlife Diseases* 43, no. 4 (October 1, 2007): 660–67. <https://doi.org/10.7589/0090-3558-43.4.660>.

Brown, Justin, Rebecca Poulson, Deborah Carter, Camille Lebarbenchon, Mary Pantin-Jackwood, Erica Spackman, Eric Shepherd, Mary Killian, and David Stallknecht. “Susceptibility of Avian Species to North American H13 Low Pathogenic Avian Influenza Viruses.” *Avian Diseases* 56, no. 4s1 (December 2012): 969–75. <https://doi.org/10.1637/10158-040912-Reg.1>.

Bublot, Michel, Alexandra Richard-Mazet, Sandrine Chanavat-Bizzini, François-Xavier Le Gros, Michelle Duboeuf, Anna Stoll, Vilmos Palfi, Eric Niqueux, Olivier Guionie, and Nick Dren. “Immunogenicíty of Poxvirus Vector Avian Influenza Vaccines in Muscovy and Pekin Ducks.” *Avian Diseases* 54, no. 1 (2010): 232–38.

Cagle, Caran, Thanh Long To, Tung Nguyen, Jamie Wasilenko, Sean C. Adams, Carol J. Cardona, Erica Spackman, David L. Suarez, and Mary J. Pantin-Jackwood. “Pekin and Muscovy Ducks Respond Differently to Vaccination with a H5N1 Highly Pathogenic Avian Influenza (HPAI) Commercial Inactivated Vaccine.” *Vaccine* 29, no. 38 (September 2, 2011): 6549–57. <https://doi.org/10.1016/j.vaccine.2011.07.004>.

Cagle, Caran, Jamie Wasilenko, Sean C. Adams, Carol J. Cardona, Thanh Long To, Tung Nguyen, Erica Spackman, et al. “Differences in Pathogenicity, Response to Vaccination, and Innate Immune Responses in Different Types of Ducks Infected with a Virulent H5N1 Highly Pathogenic Avian Influenza Virus from Vietnam.” *Avian Diseases* 56, no. 3 (September 2012): 479–87. <https://doi.org/10.1637/10030-120511-Reg.1>.

Caliendo, Valentina, Lonneke Leijten, Marco W. G. van de Bildt, Marjolein J. Poen, Adinda Kok, Theo Bestebroer, Mathilde Richard, Ron A. M. Fouchier, and Thijs Kuiken. “Long-Term Protective Effect of Serial Infections with H5N8 Highly Pathogenic Avian Influenza Virus in Wild Ducks.” *Journal of Virology* 96, no. 18 (September 13, 2022): e01233-22. <https://doi.org/10.1128/jvi.01233-22>.

Costa, T. P., J. D. Brown, E. W. Howerth, and D. E. Stallknecht. “The Effect of Age on Avian Influenza Viral Shedding in Mallards (Anas Platyrhynchos).” *Avian Diseases* 54, no. s1 (March 2010): 581–85. <https://doi.org/10.1637/8692-031309-ResNote.1>.

Costa, Taiana P., Justin D. Brown, Elizabeth W. Howerth, and David E. Stallknecht. “Variation in Viral Shedding Patterns between Different Wild Bird Species Infected Experimentally with Low-Pathogenicity Avian Influenza Viruses That Originated from Wild Birds.” *Avian Pathology* 40, no. 2 (April 1, 2011): 119–24. <https://doi.org/10.1080/03079457.2010.540002>.

Costa, Taiana P., Justin D. Brown, Elizabeth W. Howerth, David E. Stallknecht, and David E. Swayne. “Homo- and Heterosubtypic Low Pathogenic Avian Influenza Exposure on H5N1 Highly Pathogenic Avian Influenza Virus Infection in Wood Ducks (Aix Sponsa).” *PLOS ONE* 6, no. 1 (January 6, 2011): e15987. <https://doi.org/10.1371/journal.pone.0015987>.

Criado, Miria F., Kira A. Moresco, David E. Stallknecht, and David E. Swayne. “Low-Pathogenicity Influenza Viruses Replicate Differently in Laughing Gulls and Mallards.” *Influenza and Other Respiratory Viruses* 15, no. 6 (2021): 701–6. <https://doi.org/10.1111/irv.12878>.

Curran, John M., Ian D. Robertson, Trevor M. Ellis, Paul W. Selleck, and Mark A. O’Dea. “Variation in the Responses of Wild Species of Duck, Gull, and Wader to Inoculation with a Wild-Bird–Origin H6N2 Low Pathogenicity Avian Influenza Virus.” *Avian Diseases* 57, no. 3 (March 2013): 581–86. <https://doi.org/10.1637/10458-112712-Reg.1>.

Daoust, P.-Y., M. van de Bildt, D. van Riel, G. van Amerongen, T. Bestebroer, R. Vanderstichel, R. A. M. Fouchier, and T. Kuiken. “Replication of 2 Subtypes of Low-Pathogenicity Avian Influenza Virus of Duck and Gull Origins in Experimentally Infected Mallard Ducks.” *Veterinary Pathology* 50, no. 3 (May 1, 2013): 548–59. <https://doi.org/10.1177/0300985812469633>.

DeJesus, Eric, Mar Costa-Hurtado, Diane Smith, Dong-Hun Lee, Erica Spackman, Darrell R. Kapczynski, Mia Kim Torchetti, et al. “Changes in Adaptation of H5N2 Highly Pathogenic Avian Influenza H5 Clade 2.3.4.4 Viruses in Chickens and Mallards.” *Virology* 499 (December 1, 2016): 52–64. <https://doi.org/10.1016/j.virol.2016.08.036>.

Dijk, Jacintha G. B. van, Josanne H. Verhagen, Arne Hegemann, Conny Tolf, Jenny Olofsson, Josef D. Järhult, and Jonas Waldenström. “A Comparative Study of the Innate Humoral Immune Response to Avian Influenza Virus in Wild and Domestic Mallards.” *Frontiers in Microbiology* 11 (2020). <https://www.frontiersin.org/articles/10.3389/fmicb.2020.608274>.

Dolinski, Amanda C., Jared J. Homola, Mark D. Jankowski, John D. Robinson, and Jennifer C. Owen. “Differential Gene Expression Reveals Host Factors for Viral Shedding Variation in Mallards (Anas Platyrhynchos) Infected with Low-Pathogenic Avian Influenza Virus.” *The Journal of General Virology* 103, no. 3 (March 2022): 10.1099/jgv.0.001724. <https://doi.org/10.1099/jgv.0.001724>.

———. “Host Gene Expression Is Associated with Viral Shedding Magnitude in Blue-Winged Teals (Spatula Discors) Infected with Low-Path Avian Influenza Virus.” *Comparative Immunology, Microbiology and Infectious Diseases* 90–91 (November 1, 2022): 101909. <https://doi.org/10.1016/j.cimid.2022.101909>.

Dolinski, Amanda C., Mark D. Jankowski, Jeanne M. Fair, and Jennifer C. Owen. “The Association between SAα2,3Gal Occurrence Frequency and Avian Influenza Viral Load in Mallards (Anas Platyrhynchos) and Blue-Winged Teals (Spatula Discors).” *BMC Veterinary Research* 16, no. 1 (November 10, 2020): 430. <https://doi.org/10.1186/s12917-020-02642-7>.

Dolka, Beata, Artur Żbikowski, Izabella Dolka, and Piotr Szeleszczuk. “The Response of Mute Swans (Cygnus Olor, Gm. 1789) to Vaccination against Avian Influenza with an Inactivated H5N2 Vaccine.” *Acta Veterinaria Scandinavica* 58, no. 1 (October 22, 2016): 74. <https://doi.org/10.1186/s13028-016-0255-y>.

Ducatez, Mariette, Stephanie Sonnberg, Jeri Carol Crumpton, Adam Rubrum, Phouvong Phommachanh, Bounlom Douangngeun, Malik Peiris, Yi Guan, Robert Webster, and Richard Webby. “Highly Pathogenic Avian Influenza H5N1 Clade 2.3.2.1 and Clade 2.3.4 Viruses Do Not Induce a Clade-Specific Phenotype in Mallard Ducks.” *Journal of General Virology* 98, no. 6 (2017): 1232–44. <https://doi.org/10.1099/jgv.0.000806>.

Ellis, Jeremy W., J. Jeffrey Root, Loredana M. McCurdy, Kevin T. Bentler, Nicole L. Barrett, Kaci K. VanDalen, Katherine L. Dirsmith, and Susan A. Shriner. “Avian Influenza A Virus Susceptibility, Infection, Transmission, and Antibody Kinetics in European Starlings.” *PLOS Pathogens* 17, no. 8 (August 30, 2021): e1009879. <https://doi.org/10.1371/journal.ppat.1009879>.

El-Shall, Nahed A., Walaa S. H. Abd El Naby, Eid. G. S. Hussein, Ahlam E. Yonis, and Mahmoud E. Sedeik. “Pathogenicity of H5N8 Avian Influenza Virus in Chickens and in Duck Breeds and the Role of MX1 and IFN-α in Infection Outcome and Transmission to Contact Birds.” *Comparative Immunology, Microbiology and Infectious Diseases* 100 (September 1, 2023): 102039. <https://doi.org/10.1016/j.cimid.2023.102039>.

Fereidouni, Sasan R., Christian Grund, Rafaela Häuslaigner, Elke Lange, Hendrik Wilking, Timm C. Harder, Martin Beer, and Elke Starick. “Dynamics of Specific Antibody Responses Induced in Mallards After Infection by or Immunization with Low Pathogenicity Avian Influenza Viruses.” *Avian Diseases* 54, no. 1 (March 2010): 79–85. <https://doi.org/10.1637/9005-073109-Reg.1>.

Fereidouni, Sasan R., Timm C. Harder, Anja Globig, and Elke Starick. “Failure of Productive Infection of Mallards (Anas Platyrhynchos) with H16 Subtype of Avian Influenza Viruses.” *Influenza and Other Respiratory Viruses* 8, no. 6 (2014): 613–16. <https://doi.org/10.1111/irv.12275>.

Fereidouni, Sasan R., Elke Starick, Martin Beer, Hendrik Wilking, Donata Kalthoff, Christian Grund, Rafaela Häuslaigner, Angele Breithaupt, Elke Lange, and Timm C. Harder. “Highly Pathogenic Avian Influenza Virus Infection of Mallards with Homo- and Heterosubtypic Immunity Induced by Low Pathogenic Avian Influenza Viruses.” *PLOS ONE* 4, no. 8 (August 20, 2009): e6706. <https://doi.org/10.1371/journal.pone.0006706>.

França, M., R. Poulson, J. Brown, E. W. Howerth, R. D. Berghaus, D. Carter, and D. E. Stallknecht. “Effect of Different Routes of Inoculation on Infectivity and Viral Shedding of LPAI Viruses in Mallards.” *Avian Diseases* 56, no. 4s1 (December 2012): 981–85. <https://doi.org/10.1637/10151-040812-ResNote.1>.

França, M., D. E. Stallknecht, R. Poulson, J. Brown, and E. W. Howerth. “The Pathogenesis of Low Pathogenic Avian Influenza in Mallards.” *Avian Diseases* 56, no. 4s1 (December 2012): 976–80. <https://doi.org/10.1637/10153-040812-ResNote.1>.

Fujimoto, Yoshikazu, Hiroshi Ito, Kyoko Shinya, Tsuyoshi Yamaguchi, Tatsufumi Usui, Toshiyuki Murase, Hiroichi Ozaki, et al. “Susceptibility of Two Species of Wild Terrestrial Birds to Infection with a Highly Pathogenic Avian Influenza Virus of H5N1 Subtype.” *Avian Pathology* 39, no. 2 (April 1, 2010): 95–98. <https://doi.org/10.1080/03079451003599268>.

Fujimoto, Yoshikazu, Kohei Ogasawara, Norikazu Isoda, Hitoshi Hatai, Kosuke Okuya, Yukiko Watanabe, Ayato Takada, Yoshihiro Sakoda, Keisuke Saito, and Makoto Ozawa. “Experimental and Natural Infections of White-Tailed Sea Eagles (Haliaeetus Albicilla) with High Pathogenicity Avian Influenza Virus of H5 Subtype.” *Frontiers in Microbiology* 13 (2022). <https://www.frontiersin.org/articles/10.3389/fmicb.2022.1007350>.

Guionie, Olivier, Cécile Guillou-Cloarec, David Courtois, Stéphanie Bougeard, Michel Amelot, and Véronique Jestin. “Experimental Infection of Muscovy Ducks with Highly Pathogenic Avian Influenza Virus (H5N1) Belonging to Clade 2.2.” *Avian Diseases* 54, no. s1 (March 1, 2010): 538–47. <https://doi.org/10.1637/8790-040109-Reg.1>.

Gulyaeva, Marina A., Kirill A. Sharshov, Anna V. Zaykovskaia, Lidia V. Shestopalova, and Aleksander M. Shestopalov. “Experimental Infection and Pathology of Clade 2.2 H5N1 Virus in Gulls.” *Journal of Veterinary Science* 17, no. 2 (June 21, 2016): 179–88. <https://doi.org/10.4142/jvs.2016.17.2.179>.

Hall, Jeffrey S., J. Christian Franson, Robert E. Gill, Carol U. Meteyer, Joshua L. TeSlaa, Sean Nashold, Robert J. Dusek, and Hon S. Ip. “Experimental Challenge and Pathology of Highly Pathogenic Avian Influenza Virus H5N1 in Dunlin (Calidris Alpina), an Intercontinental Migrant Shorebird Species.” *Influenza and Other Respiratory Viruses* 5, no. 5 (2011): 365–72. <https://doi.org/10.1111/j.1750-2659.2011.00238.x>.

Hall, Jeffrey S., Daniel A. Grear, Scott Krauss, J. Patrick Seiler, Robert J. Dusek, Sean W. Nashold, and Robert G. Webster. “Highly Pathogenic Avian Influenza Virus H5N2 (Clade 2.3.4.4) Challenge of Mallards Age Appropriate to the 2015 Midwestern Poultry Outbreak.” *Influenza and Other Respiratory Viruses* 15, no. 6 (2021): 767–77. <https://doi.org/10.1111/irv.12886>.

Hall, Jeffrey S., Hon S. Ip, J. Christian Franson, Carol Meteyer, Sean Nashold, Joshua L. TeSlaa, John French, Patrick Redig, and Christopher Brand. “Experimental Infection of a North American Raptor, American Kestrel (Falco Sparverius), with Highly Pathogenic Avian Influenza Virus (H5N1).” *PLOS ONE* 4, no. 10 (October 22, 2009): e7555. <https://doi.org/10.1371/journal.pone.0007555>.

Hall, Jeffrey S., Scott Krauss, J. Christian Franson, Joshua L. TeSlaa, Sean W. Nashold, David E. Stallknecht, Richard J. Webby, and Robert G. Webster. “Avian Influenza in Shorebirds: Experimental Infection of Ruddy Turnstones (Arenaria Interpres) with Avian Influenza Virus.” *Influenza and Other Respiratory Viruses* 7, no. 1 (January 2013): 85–92. <https://doi.org/10.1111/j.1750-2659.2012.00358.x>.

Hall, Jeffrey S., Robin E. Russell, J. Christian Franson, Catherine Soos, Robert J. Dusek, R. Bradford Allen, Sean W. Nashold, et al. “Avian Influenza Ecology in North Atlantic Sea Ducks: Not All Ducks Are Created Equal.” *PLOS ONE* 10, no. 12 (December 17, 2015): e0144524. <https://doi.org/10.1371/journal.pone.0144524>.

Helin, Anu S., Michelle Wille, Clara Atterby, Josef D. Järhult, Jonas Waldenström, and Joanne R. Chapman. “A Rapid and Transient Innate Immune Response to Avian Influenza Infection in Mallards.” *Molecular Immunology* 95 (March 1, 2018): 64–72. <https://doi.org/10.1016/j.molimm.2018.01.012>.

Hiono, Takahiro, Masatoshi Okamatsu, Naoki Yamamoto, Kohei Ogasawara, Mayumi Endo, Saya Kuribayashi, Shintaro Shichinohe, et al. “Experimental Infection of Highly and Low Pathogenic Avian Influenza Viruses to Chickens, Ducks, Tree Sparrows, Jungle Crows, and Black Rats for the Evaluation of Their Roles in Virus Transmission.” *Veterinary Microbiology* 182 (January 15, 2016): 108–15. <https://doi.org/10.1016/j.vetmic.2015.11.009>.

Hu, J., K. Zhao, X. Liu, X. Wang, and Z. Chen. “Two Highly Pathogenic Avian Influenza H5N1 Viruses of Clade 2.3.2.1 with Similar Genetic Background but with Different Pathogenicity in Mice and Ducks.” *Transboundary and Emerging Diseases* 60, no. 2 (2013): 127–39. <https://doi.org/10.1111/j.1865-1682.2012.01325.x>.

Hu, Jiao, Yiqun Mo, Xiaoquan Wang, Min Gu, Zenglei Hu, Lei Zhong, Qiwen Wu, et al. “PA-X Decreases the Pathogenicity of Highly Pathogenic H5N1 Influenza A Virus in Avian Species by Inhibiting Virus Replication and Host Response.” *Journal of Virology* 89, no. 8 (March 19, 2015): 4126–42. <https://doi.org/10.1128/jvi.02132-14>.

Iqbal, Munir, Tahir Yaqub, Nadia Mukhtar, Muhammad Z. Shabbir, and John W. McCauley. “Infectivity and Transmissibility of H9N2 Avian Influenza Virus in Chickens and Wild Terrestrial Birds.” *Veterinary Research* 44, no. 1 (October 17, 2013): 100. <https://doi.org/10.1186/1297-9716-44-100>.

Kang, Hyun-Mi, Eun-Kyoung Lee, Byung-Min Song, Jipseol Jeong, Jun-Gu Choi, Joojin Jeong, Oun-Kyong Moon, et al. “Novel Reassortant Influenza A(H5N8) Viruses among Inoculated Domestic and Wild Ducks, South Korea, 2014.” *Emerging Infectious Diseases* 21, no. 2 (February 2015): 298–304. <https://doi.org/10.3201/eid2102.141268>.

Keawcharoen, Juthatip, Debby van Riel, Geert van Amerongen, Theo M. Bestebroer, Walter E. Beyer, Rob van Lavieren, Albert D. M. E. Osterhaus, Ron A. M. Fouchier, and Thijs Kuiken. “Wild Ducks as Long-Distance Vectors of Highly Pathogenic Avian Influenza Virus (H5N1) - Volume 14, Number 4—April 2008 - Emerging Infectious Diseases Journal - CDC,” 2008. <https://doi.org/10.3201/eid1404.071016>.

Kim, Min-Chul, Ok-Mi Jeong, Hyun-Mi Kang, Mi-Ra Paek, Ji-Sun Kwon, Chang-Seon Song, Yong-Kuk Kwon, Jung-Goo Lee, Jun-Hun Kwon, and Youn-Jeong Lee. “Pathogenicity and Transmission Studies of H7N7 Avian Influenza Virus Isolated from Feces of Magpie Origin in Chickens and Magpie.” *Veterinary Microbiology* 141, no. 3 (March 24, 2010): 268–74. <https://doi.org/10.1016/j.vetmic.2009.09.027>.

Koethe, Susanne, Lorenz Ulrich, Reiner Ulrich, Susanne Amler, Annika Graaf, Timm C. Harder, Christian Grund, et al. “Modulation of Lethal HPAIV H5N8 Clade 2.3.4.4B Infection in AIV Pre-Exposed Mallards.” *Emerging Microbes & Infections* 9, no. 1 (January 1, 2020): 180–93. <https://doi.org/10.1080/22221751.2020.1713706>.

Kwon, J.-H., D.-H. Lee, D. E. Swayne, J.-Y. Noh, S.-S. Yuk, S. Jeong, S.-H. Lee, C. Woo, J.-H. Shin, and C.-S. Song. “Experimental Infection of H5N1 and H5N8 Highly Pathogenic Avian Influenza Viruses in Northern Pintail (Anas Acuta).” *Transboundary and Emerging Diseases* 65, no. 5 (2018): 1367–71. <https://doi.org/10.1111/tbed.12872>.

Kwon, K. Y., S. J. Joh, M. C. Kim, M. S. Kang, Y. J. Lee, J. H. Kwon, and J. H. Kim. “The Susceptibility of Magpies to a Highly Pathogenic Avian Influenza Virus Subtype H5N1.” *Poultry Science* 89, no. 6 (June 1, 2010): 1156–61. <https://doi.org/10.3382/ps.2009-00549>.

Kwon, Y. K., C. Thomas, and D. E. Swayne. “Variability in Pathobiology of South Korean H5N1 High-Pathogenicity Avian Influenza Virus Infection for 5 Species of Migratory Waterfowl.” *Veterinary Pathology* 47, no. 3 (May 1, 2010): 495–506. <https://doi.org/10.1177/0300985809359602>.

Laudert, E., D. Halvorson, V. Sivanandan, and D. Shaw. “Comparative Evaluation of Tissue Trophism Characteristics in Turkeys and Mallard Ducks after Intravenous Inoculation of Type A Influenza Viruses.” *Avian Diseases* 37, no. 3 (1993): 773–80. <https://doi.org/10.2307/1592028>.

Lee, Chang-Won, and David L Suarez. “Application of Real-Time RT-PCR for the Quantitation and Competitive Replication Study of H5 and H7 Subtype Avian Influenza Virus.” *Journal of Virological Methods* 119, no. 2 (August 1, 2004): 151–58. <https://doi.org/10.1016/j.jviromet.2004.03.014>.

Leyson, Christina M., Sungsu Youk, Helena L. Ferreira, David L. Suarez, and Mary Pantin-Jackwood. “Multiple Gene Segments Are Associated with Enhanced Virulence of Clade 2.3.4.4 H5N8 Highly Pathogenic Avian Influenza Virus in Mallards.” *Journal of Virology* 95, no. 18 (August 25, 2021): 10.1128/jvi.00955-21. <https://doi.org/10.1128/jvi.00955-21>.

Leyson, Christina, Sung-su Youk, Diane Smith, Kiril Dimitrov, Dong-Hun Lee, Lars Erik Larsen, David E. Swayne, and Mary J. Pantin-Jackwood. “Pathogenicity and Genomic Changes of a 2016 European H5N8 Highly Pathogenic Avian Influenza Virus (Clade 2.3.4.4) in Experimentally Infected Mallards and Chickens.” *Virology* 537 (November 1, 2019): 172–85. <https://doi.org/10.1016/j.virol.2019.08.020>.

Li, Juan, Min Gu, Dong Liu, Benqi Liu, Kaijun Jiang, Lei Zhong, Kaituo Liu, et al. “Phylogenetic and Biological Characterization of Three K1203 (H5N8)-like Avian Influenza A Virus Reassortants in China in 2014.” *Archives of Virology* 161, no. 2 (February 1, 2016): 289–302. <https://doi.org/10.1007/s00705-015-2661-2>.

Liang, Yuan, Charlotte K. Hjulsager, Amanda H. Seekings, Caroline J. Warren, Fabian Z. X. Lean, Alejandro Núñez, Joe James, et al. “Pathogenesis and Infection Dynamics of High Pathogenicity Avian Influenza Virus (HPAIV) H5N6 (Clade 2.3.4.4b) in Pheasants and Onward Transmission to Chickens.” *Virology* 577 (December 1, 2022): 138–48. <https://doi.org/10.1016/j.virol.2022.10.009>.

Liu, Kaituo, Ruyi Gao, Min Gu, Juan Li, Liwei Shi, Wenqi Sun, Dong Liu, et al. “Genetic and Biological Characterization of Two Reassortant H5N2 Avian Influenza A Viruses Isolated from Waterfowl in China in 2016.” *Veterinary Microbiology* 224 (October 1, 2018): 8–16. <https://doi.org/10.1016/j.vetmic.2018.08.016>.

Liu, Kaituo, Min Gu, Shunlin Hu, Ruyi Gao, Juan Li, Liwei Shi, Wenqi Sun, et al. “Genetic and Biological Characterization of Three Poultry-Origin H5N6 Avian Influenza Viruses with All Internal Genes from Genotype S H9N2 Viruses.” *Archives of Virology* 163, no. 4 (April 2018): 947–60. <https://doi.org/10.1007/s00705-017-3695-4>.

Liu, Qinfang, Jingjiao Ma, Zheng Kou, Juan Pu, Fumin Lei, Tianxian Li, and Jinhua Liu. “Characterization of a Highly Pathogenic Avian Influenza H5N1 Clade 2.3.4 Virus Isolated from a Tree Sparrow.” *Virus Research* 147, no. 1 (January 1, 2010): 25–29. <https://doi.org/10.1016/j.virusres.2009.09.014>.

Luczo, Jasmina M., Diann J. Prosser, Mary J. Pantin-Jackwood, Alicia M. Berlin, and Erica Spackman. “The Pathogenesis of a North American H5N2 Clade 2.3.4.4 Group A Highly Pathogenic Avian Influenza Virus in Surf Scoters (Melanitta Perspicillata).” *BMC Veterinary Research* 16, no. 1 (September 23, 2020): 351. <https://doi.org/10.1186/s12917-020-02579-x>.

Morris, Katrina M., Anamika Mishra, Ashwin A. Raut, Eleanor R. Gaunt, Dominika Borowska, Richard I. Kuo, Bo Wang, et al. “The Molecular Basis of Differential Host Responses to Avian Influenza Viruses in Avian Species with Differing Susceptibility.” *Frontiers in Cellular and Infection Microbiology* 13 (2023). <https://www.frontiersin.org/articles/10.3389/fcimb.2023.1067993>.

Naguib, Mahmoud M., Per Eriksson, Elinor Jax, Michelle Wille, Cecilia Lindskog, Caroline Bröjer, Janina Krambrich, et al. “A Comparison of Host Responses to Infection with Wild-Type Avian Influenza Viruses in Chickens and Tufted Ducks.” *Microbiology Spectrum* 11, no. 4 (June 26, 2023): e02586-22. <https://doi.org/10.1128/spectrum.02586-22>.

Nemeth, N. M., J. D. Brown, D. E. Stallknecht, E. W. Howerth, S. H. Newman, and D. E. Swayne. “Experimental Infection of Bar-Headed Geese (Anser Indicus) and Ruddy Shelducks (Tadorna Ferruginea) With a Clade 2.3.2 H5N1 Highly Pathogenic Avian Influenza Virus.” *Veterinary Pathology* 50, no. 6 (November 1, 2013): 961–70. <https://doi.org/10.1177/0300985813490758>.

Nemeth, Nicole M., Nicholas O. Thomas, Darcy S. Orahood, Theodore D. Anderson, and Paul T. Oesterle. “Shedding and Serologic Responses Following Primary and Secondary Inoculation of House Sparrows (Passer Domesticus) and European Starlings (Sturnus Vulgaris) with Low-Pathogenicity Avian Influenza Virus.” *Avian Pathology* 39, no. 5 (October 1, 2010): 411–18. <https://doi.org/10.1080/03079457.2010.513043>.

Neufeld, J. L., C. Embury-Hyatt, Y. Berhane, L. Manning, S. Ganske, and J. Pasick. “Pathology of Highly Pathogenic Avian Influenza Virus (H5N1) Infection in Canada Geese (Branta Canadensis); Preliminary Studies.” *Veterinary Pathology* 46, no. 5 (September 1, 2009): 966–70. <https://doi.org/10.1354/vp.08-VP-0168-E-FL>.

Niqueux, Éric, Jean-Paul Picault, Michel Amelot, Chantal Allée, Josiane Lamandé, Carole Guillemoto, Isabelle Pierre, et al. “Quantitative Transmission Characteristics of Different H5 Low Pathogenic Avian Influenza Viruses in Muscovy Ducks.” *Veterinary Microbiology* 168, no. 1 (January 10, 2014): 78–87. <https://doi.org/10.1016/j.vetmic.2013.10.020>.

Nykvist, Marie, Anna Gillman, Hanna Söderström Lindström, Chaojun Tang, Ganna Fedorova, Åke Lundkvist, Neus Latorre-Margalef, Michelle Wille, and Josef D. Järhult. “In Vivo Mallard Experiments Indicate That Zanamivir Has Less Potential for Environmental Influenza A Virus Resistance Development than Oseltamivir.” *Journal of General Virology* 98, no. 12 (2017): 2937–49. <https://doi.org/10.1099/jgv.0.000977>.

Pantin-Jackwood, Mary J., Mar Costa-Hurtado, Eric Shepherd, Eric DeJesus, Diane Smith, Erica Spackman, Darrell R. Kapczynski, David L. Suarez, David E. Stallknecht, and David E. Swayne. “Pathogenicity and Transmission of H5 and H7 Highly Pathogenic Avian Influenza Viruses in Mallards.” *Journal of Virology* 90, no. 21 (October 14, 2016): 9967–82. <https://doi.org/10.1128/jvi.01165-16>.

Pantin-Jackwood, Mary J., Christopher B. Stephens, Kateri Bertran, David E. Swayne, and Erica Spackman. “The Pathogenesis of H7N8 Low and Highly Pathogenic Avian Influenza Viruses from the United States 2016 Outbreak in Chickens, Turkeys and Mallards.” *PLOS ONE* 12, no. 5 (May 8, 2017): e0177265. <https://doi.org/10.1371/journal.pone.0177265>.

Pantin-Jackwood, Mary, David E. Swayne, Diane Smith, and Eric Shepherd. “Effect of Species, Breed and Route of Virus Inoculation on the Pathogenicity of H5N1 Highly Pathogenic Influenza (HPAI) Viruses in Domestic Ducks.” *Veterinary Research* 44, no. 1 (July 22, 2013): 62. <https://doi.org/10.1186/1297-9716-44-62>.

Pasick, John, Yohannes Berhane, Carissa Embury-Hyatt, John Copps, Helen Kehler, Katherine Handel, Shawn Babiuk, et al. “Susceptibility of Canada Geese (Branta Canadensis) to Highly Pathogenic Avian Influenza Virus (H5N1) - Volume 13, Number 12—December 2007 - Emerging Infectious Diseases Journal - CDC,” 2007. <https://doi.org/10.3201/eid1312.070502>.

Pepin, K. M., K. K. VanDalen, N. L. Mooers, J. W. Ellis, H. J. Sullivan, J. J. Root, C. T. Webb, A. B. Franklin, and S. A. Shriner. “Quantification of Heterosubtypic Immunity between Avian Influenza Subtypes H3N8 and H4N6 in Multiple Avian Host Species.” *Journal of General Virology* 93, no. 12 (2012): 2575–83. <https://doi.org/10.1099/vir.0.045427-0>.

Pepin, Kim M., Clinton B. Leach, Nicole L. Barrett, Jeremy W. Ellis, Kaci K. VanDalen, Colleen T. Webb, and Susan A. Shriner. “Environmental Transmission of Influenza A Virus in Mallards.” *mBio* 14, no. 5 (September 28, 2023): e00862-23. <https://doi.org/10.1128/mbio.00862-23>.

Perkins, L. E. L., and D. E. Swayne. “Comparative Susceptibility of Selected Avian and Mammalian Species to a Hong Kong–Origin H5N1 High-Pathogenicity Avian Influenza Virus.” *Avian Diseases* 47, no. s3 (September 2003): 956–67. <https://doi.org/10.1637/0005-2086-47.s3.956>.

———. “Varied Pathogenicity of a Hong Kong-Origin H5N1 Avian Influenza Virus in Four Passerine Species and Budgerigars.” *Veterinary Pathology* 40, no. 1 (January 1, 2003): 14–24. <https://doi.org/10.1354/vp.40-1-14>.

Perkins, Laura E. Leigh, and David E. Swayne. “Susceptibility of Laughing Gulls (Larus Atricilla) to H5N1 and H5N3 Highly Pathogenic Avian Influenza Viruses.” *Avian Diseases* 46, no. 4 (October 1, 2002): 877–85. [https://doi.org/10.1637/0005-2086(2002)046[0877:SOLGLA]2.0.CO;2](https://doi.org/10.1637/0005-2086(2002)046%5b0877:SOLGLA%5d2.0.CO;2).

Phuong, Do Quy, Nguyen Tien Dung, Poul Henrik Jørgensen, Kurt Jensen Handberg, Nguyen The Vinh, and Jens Peter Christensen. “Susceptibility of Muscovy (Cairina Moschata) and Mallard Ducks (Anas Platyrhynchos) to Experimental Infections by Different Genotypes of H5N1 Avian Influenza Viruses.” *Veterinary Microbiology* 148, no. 2 (March 24, 2011): 168–74. <https://doi.org/10.1016/j.vetmic.2010.09.007>.

Ramis, Antonio, Geert van Amerongen, Marco van de Bildt, Loneke Leijten, Raphael Vanderstichel, Albert Osterhaus, and Thijs Kuiken. “Experimental Infection of Highly Pathogenic Avian Influenza Virus H5N1 in Black-Headed Gulls (Chroicocephalus Ridibundus).” *Veterinary Research* 45, no. 1 (August 19, 2014): 84. <https://doi.org/10.1186/s13567-014-0084-9>.

Reed, Mark L., Olga A. Bridges, Patrick Seiler, Jeong-Ki Kim, Hui-Ling Yen, Rachelle Salomon, Elena A. Govorkova, Robert G. Webster, and Charles J. Russell. “The pH of Activation of the Hemagglutinin Protein Regulates H5N1 Influenza Virus Pathogenicity and Transmissibility in Ducks.” *Journal of Virology* 84, no. 3 (February 2010): 1527–35. <https://doi.org/10.1128/jvi.02069-09>.

Reperant, Leslie A., Marco W. G. van de Bildt, Geert van Amerongen, Debbie M. Buehler, Albert D. M. E. Osterhaus, Susi Jenni-Eiermann, Theunis Piersma, and Thijs Kuiken. “Highly Pathogenic Avian Influenza Virus H5N1 Infection in a Long-Distance Migrant Shorebird under Migratory and Non-Migratory States.” *PLOS ONE* 6, no. 11 (November 22, 2011): e27814. <https://doi.org/10.1371/journal.pone.0027814>.

Root, J. Jeffrey, Angela M. Bosco-Lauth, Nicole L. Marlenee, and Richard A. Bowen. “Viral Shedding of Clade 2.3.4.4 H5 Highly Pathogenic Avian Influenza A Viruses by American Robins.” *Transboundary and Emerging Diseases* 65, no. 6 (2018): 1823–27. <https://doi.org/10.1111/tbed.12959>.

Sánchez-González, R., A. Ramis, M. Nofrarías, N. Wali, R. Valle, M. Pérez, A. Perlas, and N. Majó. “Infectivity and Pathobiology of H7N1 and H5N8 High Pathogenicity Avian Influenza Viruses for Pigeons (Columba Livia Var. Domestica).” *Avian Pathology* 50, no. 1 (January 2, 2021): 98–106. <https://doi.org/10.1080/03079457.2020.1832197>.

Scheibner, David, Claudia Blaurock, Thomas C. Mettenleiter, and Elsayed M. Abdelwhab. “Virulence of Three European Highly Pathogenic H7N1 and H7N7 Avian Influenza Viruses in Pekin and Muscovy Ducks.” *BMC Veterinary Research* 15, no. 1 (May 10, 2019): 142. <https://doi.org/10.1186/s12917-019-1899-4>.

Scheibner, David, Reiner Ulrich, Olanrewaju I. Fatola, Annika Graaf, Marcel Gischke, Ahmed H. Salaheldin, Timm C. Harder, Jutta Veits, Thomas C. Mettenleiter, and Elsayed M. Abdelwhab. “Variable Impact of the Hemagglutinin Polybasic Cleavage Site on Virulence and Pathogenesis of Avian Influenza H7N7 Virus in Chickens, Turkeys and Ducks.” *Scientific Reports* 9, no. 1 (August 9, 2019): 11556. <https://doi.org/10.1038/s41598-019-47938-3>.

Segovia, Karen M., Monique S. França, Christina L. Leyson, Darrell R. Kapczynski, Klaudia Chrzastek, Charlie S. Bahnson, and David E. Stallknecht. “Heterosubtypic Immunity Increases Infectious Dose Required to Infect Mallard Ducks with Influenza A Virus.” *PLOS ONE* 13, no. 4 (April 26, 2018): e0196394. <https://doi.org/10.1371/journal.pone.0196394>.

Shriner, Susan A., J. Jeffrey Root, Jeremy W. Ellis, Kevin T. Bentler, Kaci K. VanDalen, Thomas Gidlewski, and Sarah N. Bevins. “Influenza A Virus Surveillance, Infection and Antibody Persistence in Snow Geese (Anser Caerulescens).” *Transboundary and Emerging Diseases* 69, no. 2 (2022): 742–52. <https://doi.org/10.1111/tbed.14044>.

Shriner, Susan A., J. Jeffrey Root, Nicole L. Mooers, Jeremy W. Ellis, Scott R. Stopak, Heather J. Sullivan, Kaci K. VanDalen, and Alan B. Franklin. “Susceptibility of Rock Doves to Low-Pathogenic Avian Influenza A Viruses.” *Archives of Virology* 161, no. 3 (March 1, 2016): 715–20. <https://doi.org/10.1007/s00705-015-2685-7>.

Silva, Mariana Sá e, Christian Mathieu-Benson, Yong-kuk Kwon, Mary Pantin-Jackwood, and David E. Swayne. “Experimental Infection with Low and High Pathogenicity H7N3 Chilean Avian Influenza Viruses in Chiloe Wigeon (Anas Sibilatrix) and Cinnamon Teal (Anas Cyanoptera).” *Avian Diseases* 55, no. 3 (September 2011): 459–61. <https://doi.org/10.1637/9665-012011-Reg.1>.

Śmietanka, K., Z. Minta, M. Reichert, M. Olszewska, K. Wyrostek, M. Jóźwiak, and T. van den Berg. “Experimental Infection of Juvenile Domestic and Canada Geese with Two Different Clades of H5N1 High Pathogenicity Avian Influenza Virus.” *Veterinary Microbiology* 163, no. 3 (May 3, 2013): 235–41. <https://doi.org/10.1016/j.vetmic.2012.12.035>.

Soda, Kosuke, Yukiko Tomioka, Chiharu Hidaka, Mayu Matsushita, Tatsufumi Usui, and Tsuyoshi Yamaguchi. “Susceptibility of Common Family Anatidae Bird Species to Clade 2.3.4.4e H5N6 High Pathogenicity Avian Influenza Virus: An Experimental Infection Study.” *BMC Veterinary Research* 18, no. 1 (April 2, 2022): 127. <https://doi.org/10.1186/s12917-022-03222-7>.

Soda, Kosuke, Yukiko Tomioka, Tatsufumi Usui, Hiroichi Ozaki, Hiroshi Ito, Yasuko Nagai, Naoki Yamamoto, et al. “Susceptibility of Common Dabbling and Diving Duck Species to Clade 2.3.2.1 H5N1 High Pathogenicity Avian Influenza Virus: An Experimental Infection Study.” *Journal of Veterinary Medical Science* 85, no. 9 (2023): 942–49. <https://doi.org/10.1292/jvms.23-0122>.

Soda, Kosuke, Yukiko Tomioka, Tatsufumi Usui, Yukiko Uno, Yasuko Nagai, Hiroshi Ito, Takahiro Hiono, et al. “Susceptibility of Herons (Family: Ardeidae) to Clade 2.3.2.1 H5N1 Subtype High Pathogenicity Avian Influenza Virus.” *Avian Pathology* 51, no. 2 (March 4, 2022): 146–53. <https://doi.org/10.1080/03079457.2021.2022599>.

Spackman, Erica, Mary J. Pantin-Jackwood, Scott A. Lee, and Diann Prosser. “The Pathogenesis of a 2022 North American Highly Pathogenic Clade 2.3.4.4b H5N1 Avian Influenza Virus in Mallards (Anas Platyrhynchos).” *Avian Pathology* 52, no. 3 (May 4, 2023): 219–28. <https://doi.org/10.1080/03079457.2023.2196258>.

Spackman, Erica, Diann J. Prosser, Mary J. Pantin-Jackwood, Alicia M. Berlin, and Christopher B. Stephens. “THE PATHOGENESIS OF CLADE 2.3.4.4 H5 HIGHLY PATHOGENIC AVIAN INFLUENZA VIRUSES IN RUDDY DUCK (OXYURA JAMAICENSIS) AND LESSER SCAUP (AYTHYA AFFINIS).” *Journal of Wildlife Diseases* 53, no. 4 (October 1, 2017): 832–42. <https://doi.org/10.7589/2017-01-003>.

Spackman, Erica, Diann J. Prosser, Mary Pantin-Jackwood, Christopher B. Stephens, and Alicia M. Berlin. “Clade 2.3.4.4 H5 North American Highly Pathogenic Avian Influenza Viruses Infect, but Do Not Cause Clinical Signs in, American Black Ducks (Anas Rubripes).” *Avian Diseases* 63, no. 2 (January 2019): 366–70. <https://doi.org/10.1637/11950-081418-ResNote.1>.

Steensels, M., S. Van Borm, B. Lambrecht, J. De Vriese, F. -X. Le Gros, M. Bublot, and T. van den Berg. “Efficacy of an Inactivated and a Fowlpox-Vectored Vaccine in Muscovy Ducks Against an Asian H5N1 Highly Pathogenic Avian Influenza Viral Challenge (Eficacia de Una Vacuna Inactivada y Una Vacuna Recombinante Con Viruela Aviar Como Vector Contra Un $desaf\acute{i}o$ Con Una Cepa $Asi\acute{a}tica$ H5N1 de Influenza Aviar de Alta Patogenicidad En Patos Moscovitas).” *Avian Diseases* 51, no. 1 (2007): 325–31.

Stephens, Christopher B., Diann J. Prosser, Mary J. Pantin-Jackwood, Alicia M. Berlin, and Erica Spackman. “The Pathogenesis of H7 Highly Pathogenic Avian Influenza Viruses in Lesser Scaup (Aythya Affinis).” *Avian Diseases* 63, no. sp1 (December 2019): 230–34. <https://doi.org/10.1637/11909-060118-ResNote.1>.

Sun, Hailiang, Peirong Jiao, Baoqin Jia, Chenggang Xu, Liangmeng Wei, Fen Shan, Kaijian Luo, Chaoan Xin, Kouxin Zhang, and Ming Liao. “Pathogenicity in Quails and Mice of H5N1 Highly Pathogenic Avian Influenza Viruses Isolated from Ducks.” *Veterinary Microbiology* 152, no. 3 (September 28, 2011): 258–65. <https://doi.org/10.1016/j.vetmic.2011.05.009>.

Tanikawa, Taichiro, Kotaro Fujii, Yuji Sugie, Ryota Tsunekuni, Momoko Nakayama, and Sota Kobayashi. “Comparative Susceptibility of Mallard (Anas Platyrhynchos) to Infection with High Pathogenicity Avian Influenza Virus Strains (Gs/Gd Lineage) Isolated in Japan in 2004–2017.” *Veterinary Microbiology* 272 (September 1, 2022): 109496. <https://doi.org/10.1016/j.vetmic.2022.109496>.

Tanikawa, Taichiro, Saki Sakuma, Eiji Yoshida, Ryota Tsunekuni, Momoko Nakayama, and Sota Kobayashi. “Comparative Susceptibility of the Common Teal (Anas Crecca) to Infection with High Pathogenic Avian Influenza Virus Strains Isolated in Japan in 2004–2017.” *Veterinary Microbiology* 263 (December 1, 2021): 109266. <https://doi.org/10.1016/j.vetmic.2021.109266>.

Tarasiuk, Karolina, Anna Kycko, Małgorzata Knitter, Edyta Świętoń, Krzysztof Wyrostek, Katarzyna Domańska-Blicharz, Łukasz Bocian, Włodzimierz Meissner, and Krzysztof Śmietanka. “Pathogenicity of Highly Pathogenic Avian Influenza H5N8 Subtype for Herring Gulls (Larus Argentatus): Impact of Homo- and Heterosubtypic Immunity on the Outcome of Infection.” *Veterinary Research* 53, no. 1 (December 14, 2022): 108. <https://doi.org/10.1186/s13567-022-01125-x>.

Tarasiuk, Karolina, Anna Kycko, Edyta Świętoń, Łukasz Bocian, Krzysztof Wyrostek, and Krzysztof Śmietanka. “Homo- and Heterosubtypic Immunity to Low Pathogenic Avian Influenza Virus Mitigates the Clinical Outcome of Infection with Highly Pathogenic Avian Influenza H5N8 Clade 2.3.4.4.b in Captive Mallards (Anas Platyrhynchos).” *Pathogens* 12, no. 2 (February 2023): 217. <https://doi.org/10.3390/pathogens12020217>.

Tepper, Viktoria, Marie Nykvist, Anna Gillman, Erik Skog, Michelle Wille, Hanna Söderström Lindström, Chaojun Tang, Richard H. Lindberg, Åke Lundkvist, and Josef D. Järhult. “Influenza A/H4N2 Mallard Infection Experiments Further Indicate Zanamivir as Less Prone to Induce Environmental Resistance Development than Oseltamivir.” *Journal of General Virology* 101, no. 8 (2020): 816–24. <https://doi.org/10.1099/jgv.0.001369>.

Twabela, Augustin T., Masatoshi Okamatsu, Georges Mbuyi Tshilenge, Serge Mpiana, Justin Masumu, Lam Thanh Nguyen, Keita Matsuno, Isabella Monne, Bianca Zecchin, and Yoshihiro Sakoda. “Molecular, Antigenic, and Pathogenic Characterization of H5N8 Highly Pathogenic Avian Influenza Viruses Isolated in the Democratic Republic of Congo in 2017.” *Archives of Virology* 165, no. 1 (January 1, 2020): 87–96. <https://doi.org/10.1007/s00705-019-04456-x>.

Uchida, Yuko, Junki Mine, Nobuhiro Takemae, Taichiro Tanikawa, Ryota Tsunekuni, and Takehiko Saito. “Comparative Pathogenicity of H5N6 Subtype Highly Pathogenic Avian Influenza Viruses in Chicken, Pekin Duck and Muscovy Duck.” *Transboundary and Emerging Diseases* 66, no. 3 (2019): 1227–51. <https://doi.org/10.1111/tbed.13141>.

Uno, Yukiko, Kosuke Soda, Yukiko Tomioka, Toshihiro Ito, Tatsufumi Usui, and Tsuyoshi Yamaguchi. “Pathogenicity of Clade 2.3.2.1 H5N1 Highly Pathogenic Avian Influenza Virus in American Kestrel (Falco Sparverius).” *Avian Pathology* 49, no. 5 (September 2, 2020): 515–25. <https://doi.org/10.1080/03079457.2020.1787337>.

Verhagen, Josanne H., Ursula Höfle, Geert van Amerongen, Marco van de Bildt, Frank Majoor, Ron A. M. Fouchier, and Thijs Kuiken. “Long-Term Effect of Serial Infections with H13 and H16 Low-Pathogenic Avian Influenza Viruses in Black-Headed Gulls.” *Journal of Virology* 89, no. 22 (October 22, 2015): 11507–22. <https://doi.org/10.1128/jvi.01765-15>.

Verma, Asha Kumari, Manoj Kumar, Harshad V. Murugkar, Shanmugasundaram Nagarajan, Chakradhar Tosh, Pushpendra Namdeo, Rupal Singh, et al. “Experimental Infection and In-Contact Transmission of H9N2 Avian Influenza Virus in Crows.” *Pathogens* 11, no. 3 (March 2022): 304. <https://doi.org/10.3390/pathogens11030304>.

Wang, Bo, Qianqian Su, Jing Luo, Meng Li, Qiaoxing Wu, Han Chang, Juan Du, et al. “Differences in Highly Pathogenic H5N6 Avian Influenza Viral Pathogenicity and Inflammatory Response in Chickens and Ducks.” *Frontiers in Microbiology* 12 (2021). <https://www.frontiersin.org/articles/10.3389/fmicb.2021.593202>.

Wei, Liangmeng, Peirong Jiao, Yafen Song, Lan Cao, Runyu Yuan, Lang Gong, Jin Cui, et al. “Host Immune Responses of Ducks Infected with H5N1 Highly Pathogenic Avian Influenza Viruses of Different Pathogenicities.” *Veterinary Microbiology* 166, no. 3 (October 25, 2013): 386–93. <https://doi.org/10.1016/j.vetmic.2013.06.019>.

Wille, M., N. Latorre-Margalef, C. Tolf, D. E. Stallknecht, and J. Waldenström. “No Evidence for Homosubtypic Immunity of Influenza H3 in Mallards Following Vaccination in a Natural Experimental System.” *Molecular Ecology* 26, no. 5 (March 2017): 1420–31. <https://doi.org/10.1111/mec.13967>.

Wille, Michelle, Caroline Bröjer, Åke Lundkvist, and Josef D. Järhult. “Alternate Routes of Influenza A Virus Infection in Mallard (Anas Platyrhynchos).” *Veterinary Research* 49, no. 1 (October 29, 2018): 110. <https://doi.org/10.1186/s13567-018-0604-0>.

Yamamoto, Yu, Kikuyasu Nakamura, Manabu Yamada, and Masaji Mase. “Limited Susceptibility of Pigeons Experimentally Inoculated with H5N1 Highly Pathogenic Avian Influenza Viruses.” *Journal of Veterinary Medical Science* 74, no. 2 (2012): 205–8. <https://doi.org/10.1292/jvms.11-0312>.

———. “Pathogenesis in Eurasian Tree Sparrows Inoculated with H5N1 Highly Pathogenic Avian Influenza Virus and Experimental Virus Transmission from Tree Sparrows to Chickens.” *Avian Diseases* 57, no. 2 (January 25, 2013): 205–13. <https://doi.org/10.1637/10415-101012-Reg.1>.

Youk, Sung-Su, Dong-Hun Lee, Christina M. Leyson, Diane Smith, Miria Ferreira Criado, Eric DeJesus, David E. Swayne, and Mary J. Pantin-Jackwood. “Loss of Fitness of Mexican H7N3 Highly Pathogenic Avian Influenza Virus in Mallards after Circulating in Chickens.” *Journal of Virology* 93, no. 14 (June 28, 2019): 10.1128/jvi.00543-19. <https://doi.org/10.1128/jvi.00543-19>.

This work was funded by the U.S. Geological Survey, Ecosystem Mission Area. All data reported in this manuscript are publicly available. Any use of trade, firm, or product names is for descriptive purposes only and does not imply endorsement by the U.S. Government.
